# Frame Editors for Precise, Template-Free Frameshifting

**DOI:** 10.1101/2022.12.05.518807

**Authors:** Shota Nakade, Kazuki Nakamae, Tzu-Chieh Tang, Dou Yu, Tetsushi Sakuma, Takashi Yamamoto, Timothy K. Lu

## Abstract

Efficiency and accuracy are paramount in genome editing. While CRISPR-Cas nucleases are efficient at editing target genes, their accuracy is limited because following DNA cleavage by Cas proteins, error-prone repair mechanisms introduce random mutations. Improving the accuracy of CRISPR-Cas by reducing random repairs using DNA- or RNA-based templates can compromise efficiency. To simultaneously improve both editing efficiency and accuracy, we created a frameshifting genome-editing technology by fusing Cas9 with DNA polymerases. These Frame Editors (FEs) introduce precise and controlled frameshifts into target loci via specific DNA repairs near Cas9-induced cleavage loci. We demonstrate two types of FEs: the insertion-inducing frame editor (iFE) and the deletion-inducing frame editor (dFE). For iFE, DNA polymerase beta (POLB) is fused with Cas9, which increases the frequency of 1-bp insertions. For dFE, T4 DNA polymerase (T4pol) is fused with Cas9, which increases the frequency of 1-bp deletions. Both types of FEs reduce the number of random mutations at target loci compared with Cas9. We show that off-target editing can be reduced by substituting Cas9 with high-fidelity variants, such as HiFi Cas9 or LZ3 Cas9. Thus, FEs can introduce frameshifts into target loci with much improved mutation profiles compared with Cas9 alone and without the requirement for template sequences, offering a new strategy for repairing pathogenic frameshifts.

## Introduction

Highly efficient gene editing is fundamental to advancing biological research and developing new genetic medicines. The clustered regularly interspaced short palindromic repeats (CRISPR)-Cas9 system provides versatile ways to edit genetic information^1–4^ because it can easily target specific genomic sequences for modification. This system was originally based on the adaptive immune system of bacteria^5^, in which a CRISPR-associated nuclease, such as Cas9, prevents the proliferation and propagation of foreign genetic elements by forming an effector complex with target-recognizing CRISPR RNA (crRNA), and inducing sequence-specific cleavage^6^.

CRISPR genome editing utilizes a single guide RNA (sgRNA) and a Cas protein. The sgRNA recognizes the target region of the genome via the spacer sequence, and the nuclease domain of the Cas protein introduces a double-strand break (DSB) into the recognized sequence. There are several routes to repairing DSBs. When the DSBs are repaired by error-prone repair pathways, insertions and deletions (indels) may be introduced. Non-homologous end joining (NHEJ), which is the dominant repair pathway that can work through the cell cycle^7^, directly ligates the DSB ends and often introduces small changes at the junction. Microhomology-mediated end joining (MMEJ) is also responsible for some mutations introduced by Cas9^8^. MMEJ generates a DNA end resection by exonuclease activity, and micro-homologies (MHs) then anneal around the cleavage site, resulting in deletions^9^. Both NHEJ and MMEJ work in various cells and organisms to create Cas-induced indels that disrupt the function of targeted genes^10, 11^. However, these repair mechanisms are not suitable for implementing precise genetic modifications, such as specific frameshifts or knockouts.

It is known that Cas9-induced mutations are not completely random and are biased. Recent reports have predicted the results of NHEJ/MMEJ-mediated gene editing using large-scale mutation measurements^12–15^. These reports revealed that 1-base pair (bp) insertions and precise deletions can result from MMEJ, whereas small deletions can occur independently of MHs.

1-bp insertions constitute one particularly biased type of mutations among those introduced by CRISPR-Cas9. NHEJ can fill in overhangs mediated by CRISPR-Cas9 and introduce 1-bp insertions^13,15–17^. Specifically, double-strand cleavage is carried out by the two nuclease domains, HNH and RuvC, of Cas9. Molecular dynamics simulation showed that the HNH domain of SpCas9 cleaves target DNA 3-bp away from the protospacer-adjacent motif (PAM) sequence, whereas the RuvC domain may work closer to 4-bp than to 3-bp^18^. The 1-bp staggered ends generated by SpCas9 can be filled in with NHEJ-related enzymes such as the X-family of DNA polymerases, resulting in 1-bp insertions^16,17,19^.

Algorithms based on these insights can assist in predicting sgRNAs with desired functionalities for gene disruption and treating pathogenic mutations^12–15^. These prediction tools can be a powerful means of enabling accurate and template-free gene modifications by CRISPR-Cas9. However, there is still a significant need to enhance the efficiencies of specific types of mutations that can result from CRISPR genome editing.

There have been several reported ways to control editing outcomes without using a donor template^20–24^. One approach is to manipulate the DSB repair process, which is exemplified in the work undertaken by Sherwood *et al*.^20^, who showed that an ATM kinase inhibitor could increase the frequency of 1-bp insertion outcomes with genome editing in human and mouse cells^20^. Although this study represented a significant advance in using a small molecule to control Cas9 editing outcomes, the drawback is that additional steps, such as drug administration or gene disruption, are needed to regulate the DSB repair process. Another way to control editing outcomes is to modify the end cleaved by Cas9^21,22^. Zhang *et al*. designed the LZ3 Cas9 variant, which is a high-specificity variant of SpCas9 that increases the frequency of 1-bp insertions^21^. When there is a guanine at position 2-bp 5′ to the PAM, LZ3 Cas9 increases the frequency of 1-bp insertions compared to wild-type SpCas9. This increase in 1-bp insertion frequency is probably due to the change in the form of the cleavage end by LZ3 Cas9 under the above conditions. The application of LZ3 Cas9 suggests that CRISPR editing outcomes can be modified by simply changing a cleavage pattern.

To increase the range of applications for genome editing, we sought to create enhanced genome editors to efficiently induce 1-bp indels. Such genome editors would enable more effective gene knockouts or the ability to correct pathogenic frameshifts. To do this, we created frame editors (FEs) by combining Cas9 with an enzyme that processes the cleavage ends. FEs consist of two fused parts: CRISPR-Cas9 and DSB-end processing polymerases. One of these polymerases is human DNA polymerase beta (POLB)^25^, which is the smallest human polymerase and has only a single Pol-β-like domain, a commonly conserved feature in family X polymerases. Another polymerase that we tested was phage-derived T4 DNA polymerase (T4pol)^26^, which blunts overhangs created by restriction enzymes; because it has both polymerase and exonuclease activities^27^, it was thus expected to produce different editing results from those of POLB.

We hypothesize that when Cas9 generates 1-bp staggered cut ends, the polymerases fill in the 1-bp overhangs and precisely link those ends by NHEJ to introduce a 1-bp insertion (Figure 1a). Consequently, we created an FE that could induce 1-bp insertions and another FE that could generate 1-bp deletions. These FEs further enhance our ability to precisely control the outcome of CRISPR-Cas9 gene editing.

**Figure 1.**
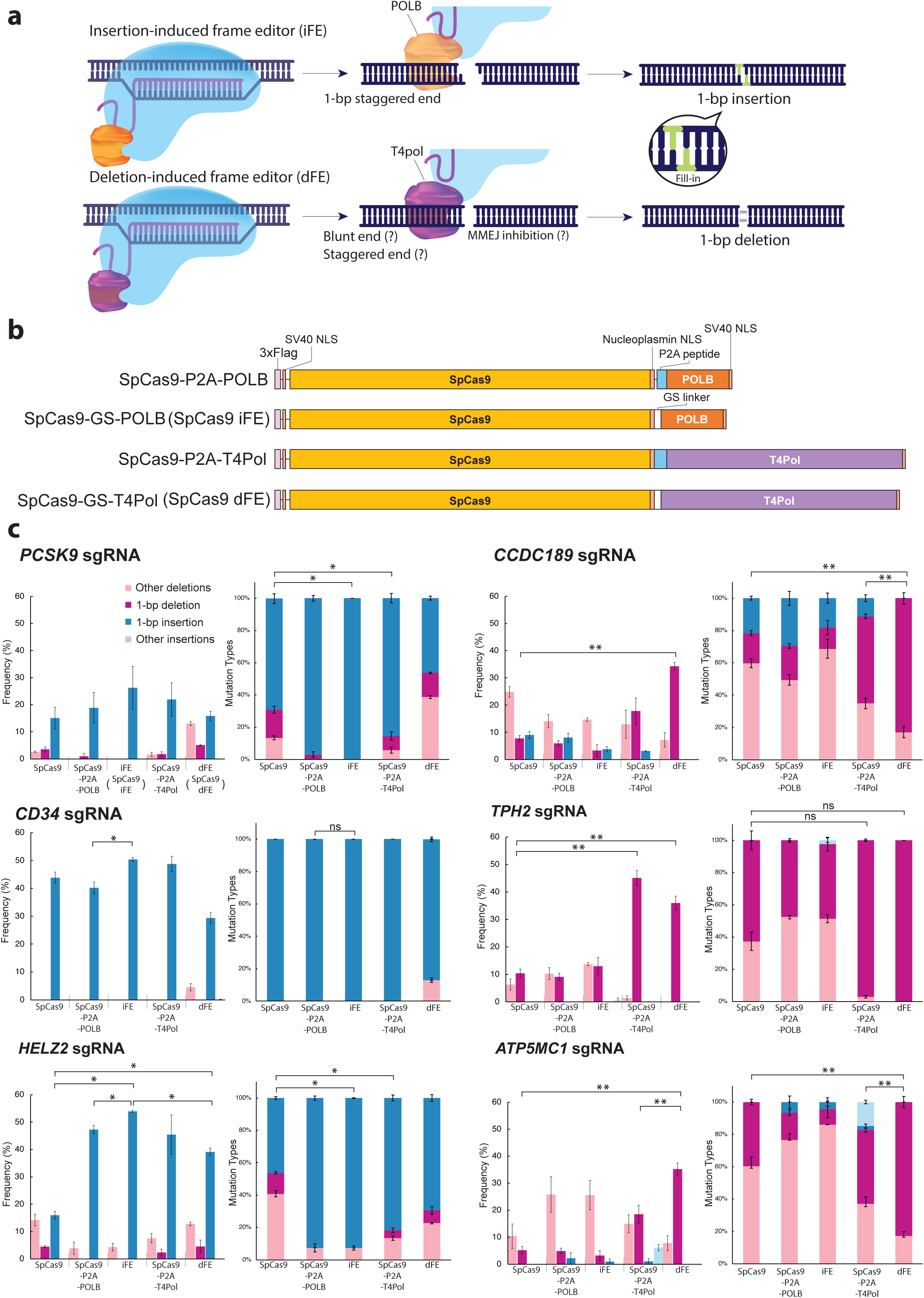
Design of frame editors (FEs) and feasibility studies. **a**. Schematic diagram showing a 1-bp indel induced by a DSB-end processing enzyme. CRISPR-Cas9 cleaves the target sequence, leaving either blunt or 1-bp staggered ends. Once staggered ends are generated, DSB-end processing enzymes process them by introducing 1-bp indels. **b**. Structure of FEs. A DSB-end processing polymerase is fused to the C-terminal end of SpCas9 with a (GGGS)_3_ linker. We compared the fusion designs to co-expression of SpCas9 with DSB-end processing polymerases with an intervening P2A self-cleaving peptide. **c**. Mutation frequencies of SpCas9, SpCas9 FEs, SpCas9-P2A-POLB, and SpCas9-P2A-T4pol were measured by TIDE analysis in HEK293T cells. Shown for each sgRNA are the frequency of mutation types in sequence traces (left) and the frequency of mutation types among total mutations (right). TIDE analysis was calculated by setting the indel size range at 1 to 10 and excluding those with P ≥ 0.001. Mean ± s.e.m. of n = 3 independent biological replicates. 1-bp insertion: *P<0.05, 1-bp deletion: **P<0.05 (*Welch’s t-test*).

## Results

### Construction of frame editors that regulate Cas9-mediated editing outcomes

We first confirmed that DSB-end processing polymerases alter editing results during gene editing, and also simultaneously investigated whether the polymerases functioned more effectively if they were bound to SpCas9 or if they were independently expressed (Figure 1). To do this, we created Cas9-polymerase fusions using a (GGGS)_3_ linker (GS hereafter), or co-expressed SpCas9 and the polymerase from a single transcript using an intervening P2A sequence, which causes co-translational cleavage of the encoded polypeptide (Figure 1b). Select sgRNAs targeting *PCSK9, CD34, HELZ2, CCDC189, TPH2, ATP5MC1, XRCC6, and GGACT* loci were used for this experiment (Supplementary Table 1). Ultimately, we found that fusing SpCas9 to polymerases resulted in efficient one-component systems that could introduce 1-bp indels (Figure 1c, S1a).

Of the bound and unbound polymerases we tested, SpCas9-GS-POLB was the most efficient at introducing 1-bp insertions (Figure 1c, S1a). We named this fusion protein the “SpCas9 insertion-inducing frame editor” (SpCas9 iFE). The SpCas9 iFE introduced an increased fraction of 1-bp insertions, and a corresponding reduction in all other mutation types, at the *PCSK9* and *HELZ2* target loci.

SpCas9-GS-T4pol was designated as a deletion-inducing frame editor (SpCas9 dFE) because it increased the frequency of 1-bp deletions (Figure 1c, S1a). This construct generated frequent 1-bp deletions and reduced other mutation types at the *CCDC189 (also known as CFAP119), TPH2, ATP5MC1*, and *GGACT* loci. For the sgRNAs targeting the *HELZ2* locus, the SpCas9 dFE also increased the number of 1-bp insertions, but the number of 1-bp insertions at this locus was higher with the SpCas9 iFE.

### Characterization of mutations induced by frame editors

To more precisely characterize mutational patterns introduced by SpCas9 iFE and SpCas9 dFE, we used 20 different sgRNAs to target various loci in HEK293T cells (Supplementary Table 1). In order to measure changes in editing frequency, a GFP expression plasmid was introduced simultaneously with the plasmid for genome editing, GFP-positive cells were sorted before genome extraction, and then sequencing data was collected by next-generation sequencing (NGS).

For five sgRNAs, SpCas9 iFE increased the frequency of 1-bp insertions significantly, by 1.3- to 2.4-fold (1.8-fold on average), compared with mutations caused by SpCas9 paired with the same sgRNAs, and decreased other mutation frequencies (Figure 2a, b, S1b). For those mutations containing 1-bp insertions, the inserted base was almost always the same as the base located 4-bp upstream from the PAM site (Figure 2d, S4a). When cytosine was located 4-bp upstream of the PAM site, SpCas9 iFE editing greatly increased the frequency of 1-bp cytosine insertions compared with the SpCas9 control. These results support our hypothesis that POLB introduces a 1-bp insertion by filling in the 1-bp overhang generated by SpCas9 because SpCas9 endonuclease cleaves 4 to 3 nt upstream from the 3’ end of the spacer region to generate a 1-bp overhang.

**Figure 2.**
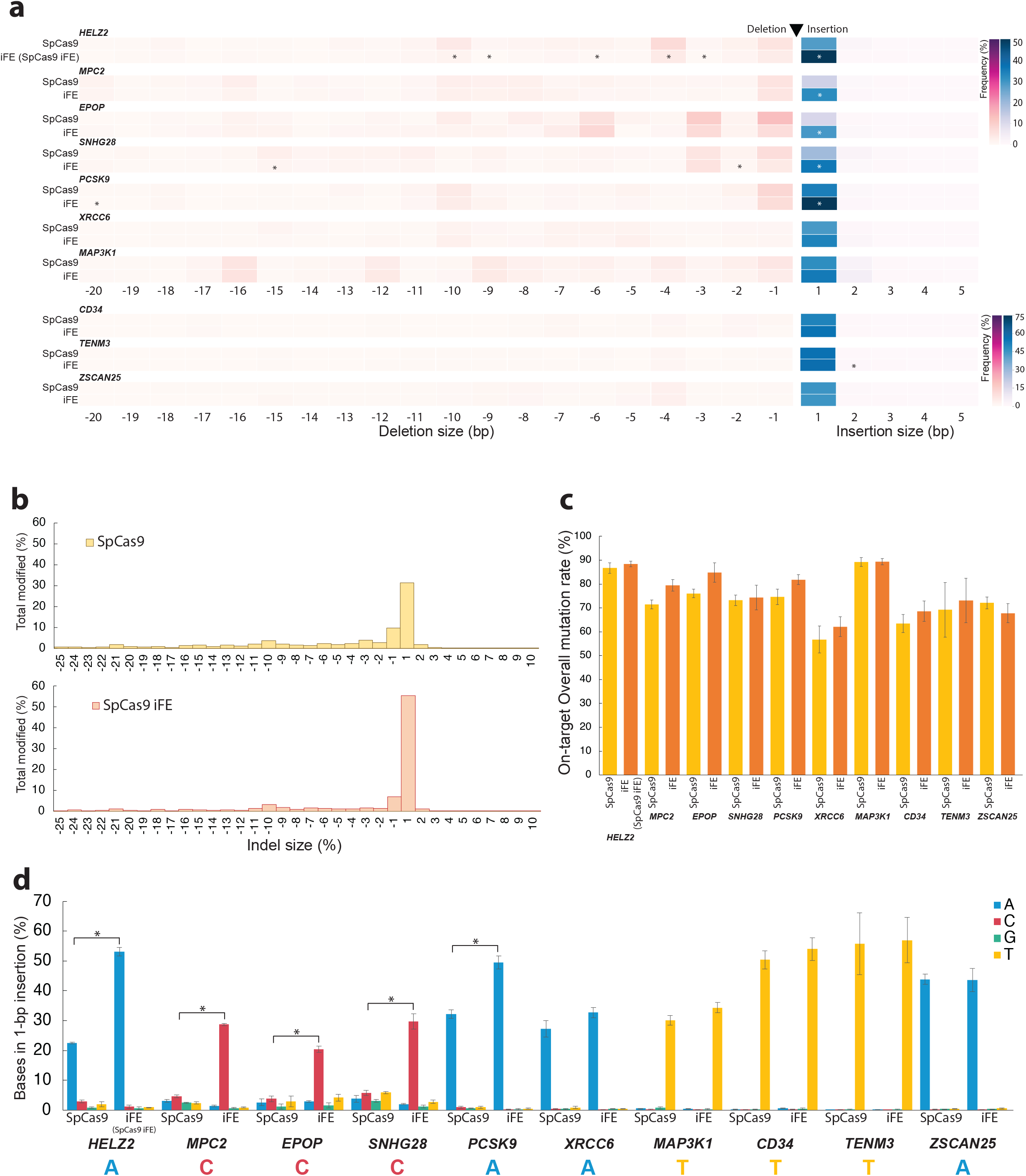
Insertion-inducing frame editor (iFE) increases the frequencies of 1-bp insertions. Comparison of mutation types induced by SpCas9 and SpCas9 iFE in HEK293T cells. **a**. Distribution of indel size. The heatmap indicates the frequency of insertion (blue) and deletion (red) in total reads. Target genes of sgRNAs are shown in the upper left. SpCas9 (top) and SpCas9 iFE (bottom) are shown on the y-axis for each sgRNA; the x-axis shows indel size. **b**. Total percentages by indel length of SpCas9 and SpCas9 iFE in all targets with significant differences. **c**. Frequencies of on-target overall mutation. **d**. Frequency of inserted bases in 1-bp insertions. The bases on staggered ends are shown at the bottom of the x-axis. Only transfected cells were collected by GFP+ sorting. Data are expressed as means□±□s.e.m. of n = 3 biological replicates. *P<0.05 (*Welch’s t-test*).

For half of the sgRNAs tested, the SpCas9 dFE significantly increased 1-bp deletions. The frequency of 1-bp deletion increased significantly, by 1.7- to 4.1-fold (2.6-fold on average), compared to that seen with SpCas9, and there were fewer other mutation types (Figure 3a, b, S1c). With 7 sgRNAs, SpCas9 dFE induced fewer in-frame deletions, which are multiples of 3-bp in length (Figure 3d). The frequency of non-MH-associated deletions increased significantly at all target loci whereas in-frame mutations decreased (Figure 3d, 3e, S4c). This effect suggested that SpCas9 dFE also suppresses deletions by MMEJ. It has been reported that for some sgRNAs, in-frame mutations generated through MMEJ repair may constitute a large percentage of the mutations induced by conventional CRISPR genome editing^8^, resulting in incomplete disruption of protein function. FEs would suppress these incomplete mutations and generate more effective protein knockouts.

**Figure 3.**
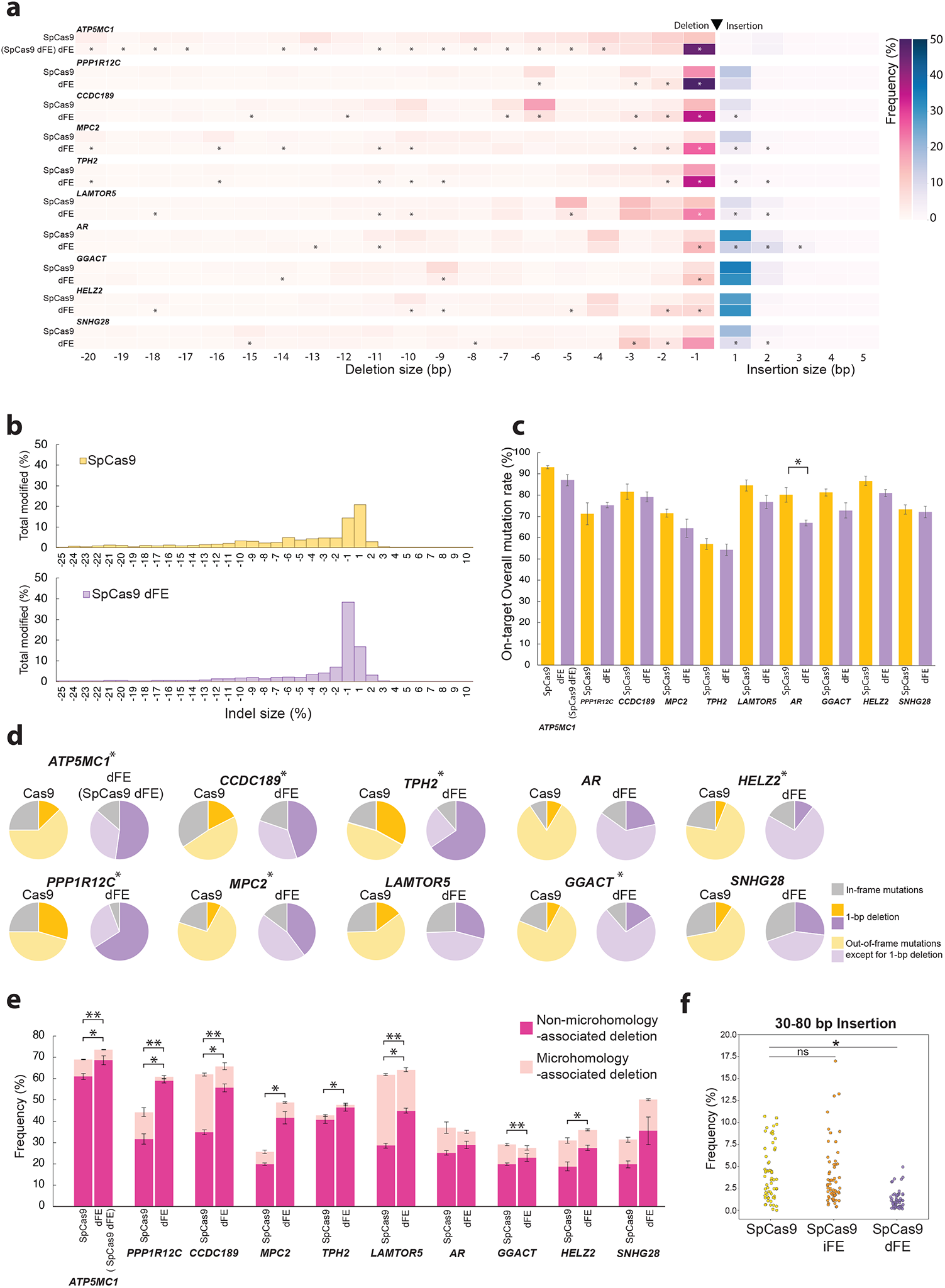
Deletion-inducing frame editor (dFE) increases the frequencies of 1-bp deletions. Comparison of mutations induced by SpCas9 and SpCas9 dFE in HEK293T cells. **a**. Distribution of indel size. Target genes of sgRNA are shown in the upper left. The heatmap indicates the frequency of insertion (blue) and deletion (red) in total reads. Shown for each sgRNA are SpCas9 (top) and SpCas9 dFE (bottom) on the y-axis, and indel size on the x-axis. **b**. Total percentages by indel length in all targets having significant differences. **c**. Frequencies of on-target overall mutation. **d**. Frequencies of 1-bp deletion, out-of-frame (deletion length is not a multiple of 3, except for a 1-bp deletion) and in-frame mutations (deletion length is multiple of 3) in on-target overall mutation. Out-of-frame and in-frame mutation is the sum of all mutations with indel lengths in each condition. **e**. Frequencies of deletions associated with non-microhomologies or microhomologies. Mutation reads with deletion lengths 15-bp or less were analyzed, and those with ≥2-bp homology were annotated as microhomologies. Only transfected cells were collected by GFP+ sorting. Non-microhomology-associated deletions: *P<0.05, Microhomology-associated deletions: **P<0.05 (*Welch’s t-test*). **f**. Comparison of large-size insertions induced by SpCas9, SpCas9 iFE, and SpCas9 dFE in HEK293T cells. Total frequencies of 30−80 nt indels for each sgRNA. Only transfected cells were collected by GFP+ sorting. Data are expressed as means□±□s.e.m. of n = 3 biological replicates. *P<0.05 (*Welch’s t-test*).

Importantly, we also found that SpCas9 dFE decreased the number of insertions 30–80 nt long by up to 7.3% (Figure 3f, S2a), while deletions 30-80 nt long remained unchanged (Figure S2b, S2c). Unexpected large insertions have been reported for genome editing by SpCas9, in which the inserted nucleotides were derived from various origins, such as genomic DNA, mRNA, exogenous vectors, or exosomes^28,29^. This data suggests that SpCas9 dFE could be used to suppress these unexpected long insertions.

The overall on-target mutation rate by iFE and dFE did not change much compared to those observed with SpCas9 (Figure 2c, 3c). The on-target overall mutation rate by SpCas9 and the SpCas9 FEs seemed to depend on the cleavage efficiency of Cas nuclease, rather than the processing of cleavage ends.

We also compared the differences between inDelphi model predictions and experimental outcomes for SpCas9 iFE and dFE using Kullback-Leibler divergence as a measure to determine the extent to which SpCas9 FEs alter the distribution of mutation patterns by SpCas9 (Figure S3). This analysis showed that the distribution of mutation patterns induced by SpCas9 dFE differed dramatically from those by SpCas9. On the other hand, SpCas9 iFE had a similar divergence score as SpCas9, with the exception of the sgRNA targeting the *HELZ2* locus. SpCas9 iFE primarily increased the frequency of 1-bp insertions, while SpCas9 dFE mainly increased the frequency of 1-bp deletions (Figure S1b, c, S4a, b). Thus, FEs can be used to introduce frameshifts in targets by skewing the distribution of mutations towards single bp insertions or deletions.

### Frame editors using HiFi Cas9 and LZ3 Cas9

Because the overall on-target mutation frequency introduced by FEs is similar to that of Cas9, we predicted that off-target effects would also be similar. We first verified that the FEs could be constructed using HiFi Cas9, a variant of Cas9 with a R691A mutation that augments both activity and specificity^30^. We created HiFi FEs by joining HiFi Cas9 either with POLB (HiFi iFE) or with T4pol (HiFi dFE). We then examined the off-target effects of the SpCas9 FEs and HiFi FEs.

In HEK293T cells, HiFi dFE increased the frequency of 1-bp deletions significantly at *CCDC189, ATP5MC1, TPH2, MPC2, GGACT, PPP1R12C*, and *PCSK9* locI (Figure 4a), by 1.6-to 5.6-fold compared with HiFi Cas9. Few other deletion types were observed with HiFi dFE. With sgRNAs targeting the *ATP5MC1, PPP1R12C*, and *PCSK9* loci, the frequencies of 1-bp deletions with HiFi dFE were lower than those observed with SpCas9 dFE (Figure 4b). In contrast, HiFi iFE did not induce a significant number of 1-bp insertions with the sgRNAs we used compared to HiFi Cas9 (Figure 4a).

**Figure 4.**
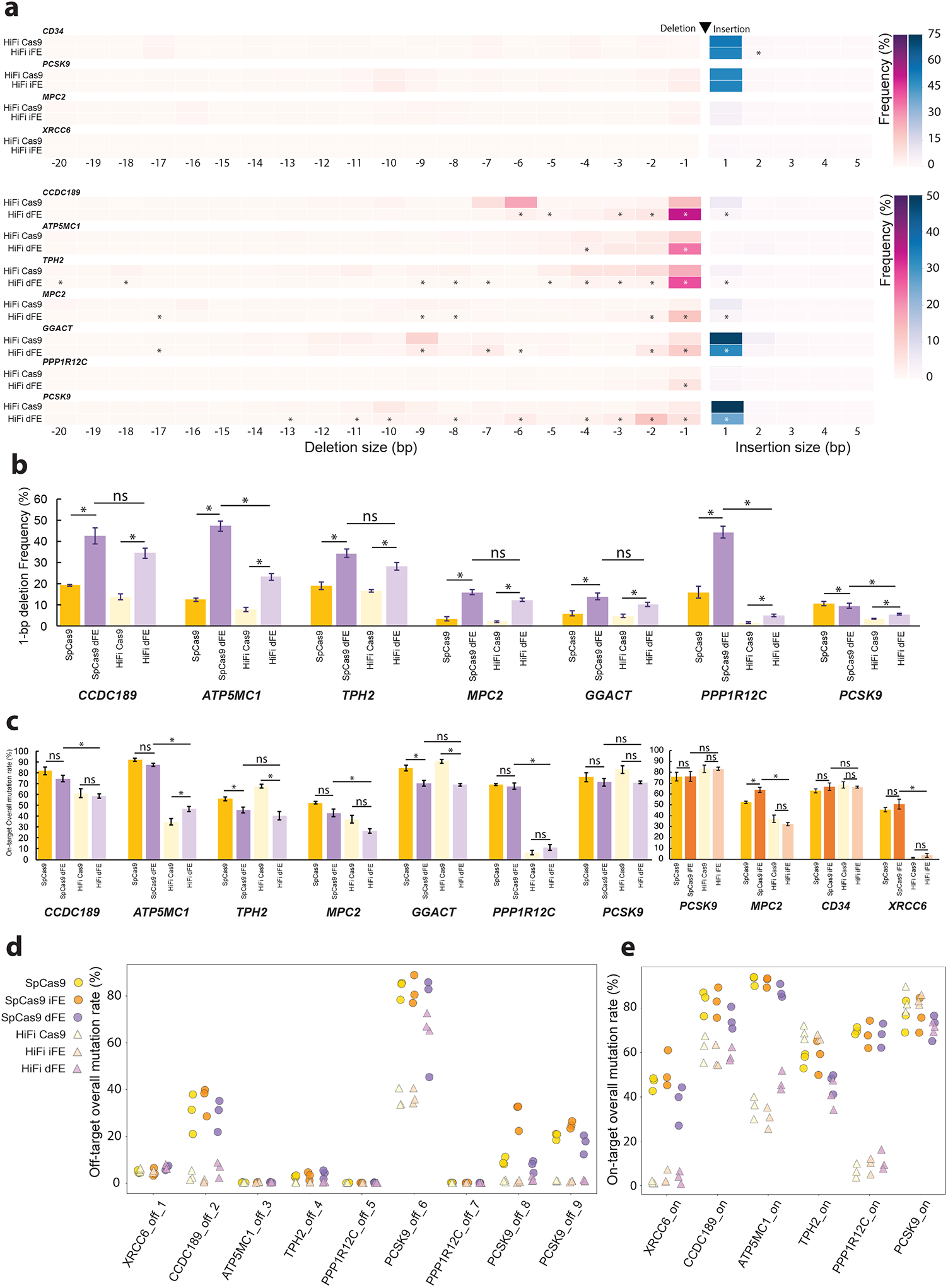
Off-target effects and FE using HiFi Cas9 (HiFi FE). **a-c**. Editing outcomes of HiFi FEs shown as heatmap by indel size **(a)**, efficiencies of 1-bp deletions compared with SpCas9 FEs **(b)**, and on-target overall mutation rates **(c)**. Data are expressed as mean□±□s.e.m. of n = 3 biological replicates. *P<0.05 (*Welch’s t-test*). **d-e**. Off-target **(d)** and On-target effects **(e)** of SpCas9, SpCas9 FEs, HiFi Cas9, and HiFi FEs. Genomic loci with high possibilities of off-target were identified using MIT scores among all sgRNAs. Only transfected HEK293T cells were collected by GFP+ sorting.

Compared with those generated by SpCas9 and SpCas9 FEs respectively, overall on-target mutation rates were decreased by HiFi Cas9 and HiFi FEs for the sgRNAs targeting the *CCDC189, ATP5MC1, MPC2, PPP1R12C*, and *XRCC6* loci (Figure 4c). This suggests that the on-target mutation activities of some the sgRNAs were reduced by HiFi Cas9 compared with those by SpCas9.

In addition, we examined on-target mutation patterns using the LZ3 Cas9 variant by building LZ3 iFE and LZ3 dFE, which preferentially induced 1-bp indels at the target loci when used with sgRNAs containing a guanine 2-bp upstream of PAM site^21^ (Figure S5). With 5 of the 8 sgRNAs tested, LZ3 FEs increased the frequency of 1-bp indels compared with LZ3 Cas9 (Figure S6). Among those sgRNAs with a G at -2 PAM (the *PCSK9, MPC2, MAP3K1*, and *XRCC6* loci), LZ3 iFE significantly increased the number of 1-bp insertions at the *PCSK9* and *MPC2* loci, compared with the LZ3 Cas9. The sgRNA targeting the *MPC2* locus with a G at -2 PAM mostly induced 1-bp insertions with LZ3 Cas9, whereas SpCas9 introduced numerous other indel types in addition. Interestingly, LZ3 dFE increased the frequency of 1-bp insertion at the *MPC2* locus, whereas SpCas9 dFE locus increased the frequency of 1-bp deletion. In contrast, for sgRNAs without G at -2 PAM, LZ3 dFE significantly increased 1-bp deletions.

### Off-target effects by frame editors

We searched for potential off-target editing sites by FEs using the CRISPOR tool (http://crispor.tefor.net/)^31^. In this scoring algorithm, the higher the MIT off-target score, the higher the risk of introducing an off-target mutation. Off-target sites were calculated for 20 sgRNAs using NGS analysis, and the 9 sites with the high MIT off-target scores were selected as the off-target sites for frame editing (Figure 4d, Supplementary Table 2).

The frequencies of the overall off-target mutations induced by SpCas9 iFE and SpCas9 dFE at eight of these loci differed only by ±5% on average from that induced by SpCas9. Only one site, PCSK9_off_8, had a 20% higher off-target overall mutation frequency induced by the SpCas9 iFEs. At sites ATP5MC1_off_3, PPP1R12C_off_5, and PPP1R12C_off_7, where the overall off-target mutations generated by SpCas9 were very low, those caused by SpCas9 FEs were also low. This observation provides evidence that the off-targeting effects of the SpCas9 FEs depend on the cleavage activity of the SpCas9 nuclease.

With HiFi Cas9 and HiFi FEs, the frequencies of overall off-target mutations at 8 of the 9 sites (sites 1–5 and 7–9) nearly diminished to zero. Site PCSK9_off_6 displayed an unusually high frequency of off-target editing by all tools. Even so, HiFi Cas9 and HiFi iFE reduced the frequency of overall off-target mutations compared with SpCas9 and SpCas9 FEs. However, HiFi dFE increased the overall off-target mutation rates at site PCSK9_off_6, similar to SpCas9 and SpCas9 FEs. We found that SpCas9 iFE increased the number of off-target 1-bp insertions compared with SpCas9 at sites CCDC189_off_2, PCSK9_off_8 and PCSK9_off_9 (Figure S6). SpCas9 dFE increased the frequency of off-target 1-bp deletions when compared with SpCas9 at sites CCDC189_off_2, TPH2_off_4, and PCSK9_off_6. For targets such as *CCDC189, ATP5MC1*, and *TPH2*, it was effective to use HiFi FEs due to low off-target editing with concomitant efficient on-target editing (Figure 4e).

The overall off-target mutation frequencies by LZ3 Cas9 and LZ3 FEs were analyzed using off-target sites CCDC189_off_2, PCSK9_off_6, and PCSK9_off_9, which had enough off-target mutations to be identified by Sanger sequencing (Figure S7). LZ3 Cas9 and the LZ3 FEs decreased off-target effects at all sites compared with SpCas9 and SpCas9 Fes, except for LZ3 dFE at site PCSK9_off_6.

We thus conclude that for sgRNAs with a high risk of off-target mutations (e.g., an MIT score greater than 2), it would be beneficial to use HiFi FEs or LZ3 FEs rather than SpCas9 FEs.

### sgRNA prediction available for frame editors

We showed that FEs robustly increase 1-bp frame editing, but the impact is sgRNA-dependent (Figure 2 and 3). To establish a strategy for selecting optimal sgRNAs, we sought to repurpose the established sgRNA prediction tool inDelphi^14^, which is intended for SpCas9, to identify sgRNAs for SpCas9 FEs.

We first experimentally confirmed that the SpCas9-mediated mutation types correlated with predicted mutation types by inDelphi in a sgRNA-specific manner (Figure 5a, b) (R-squared coefficient of determination = 0.64 for 1-bp insertions; 0.69 for 1 - 59 bp total deletions). Based on this prediction platform, we compared the mutation types mediated by SpCas9 FEs using the same sgRNAs.

**Figure 5.**
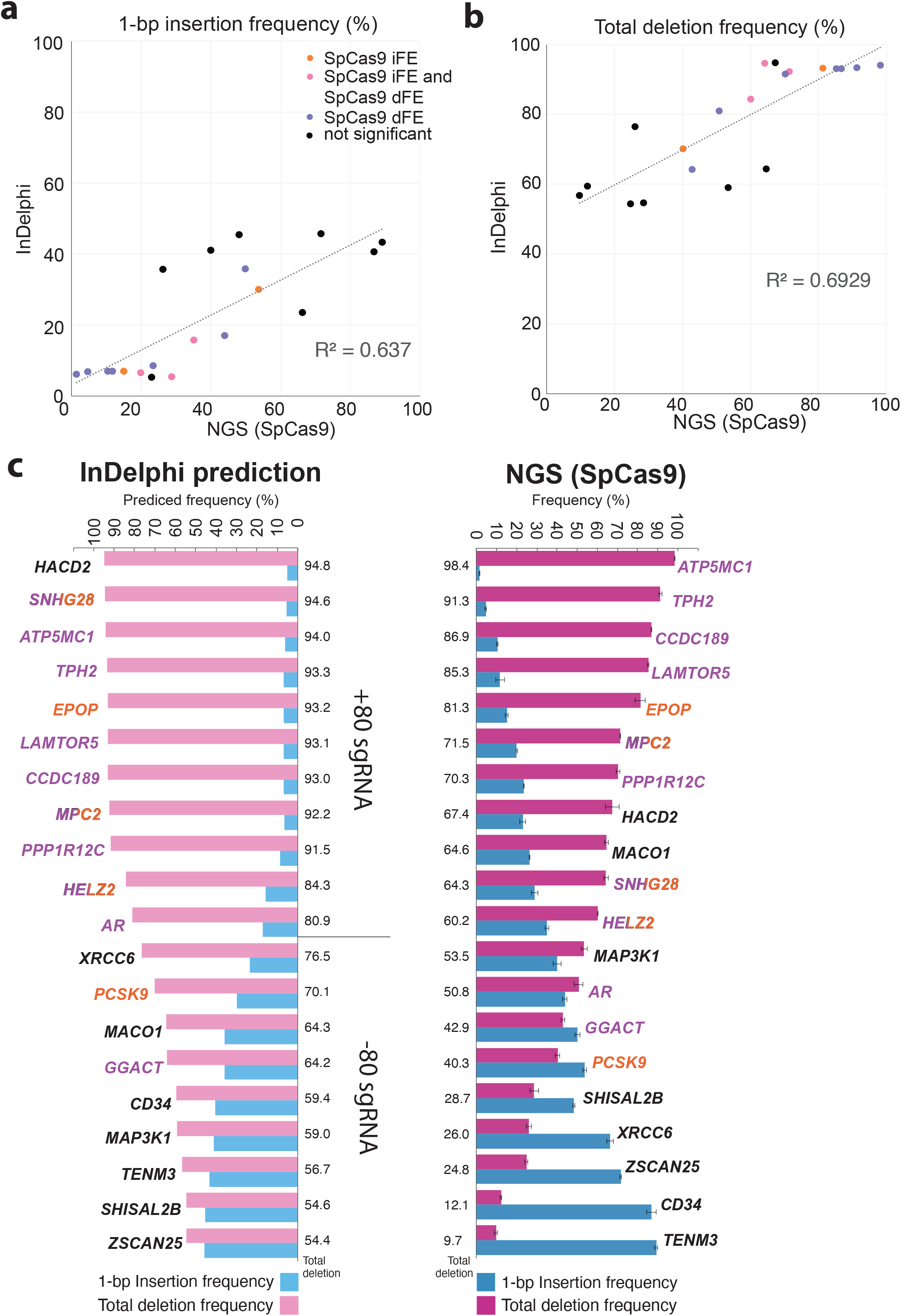
Comparison of inDelphi predictions and NGS outcomes. **a-b**. Total frequencies of 1-bp insertions **(a)** and total deletions **(b)** calculated from inDelphi predictions (y-axis) compared to NGS-measured experimental outcomes (x-axis). Correlations between inDelphi and NGS measurements were calculated by linear regression (solid line), with Pearson correlation coefficients (R) displayed. The NGS mutation outcomes by SpCas9 were also limited to those of the same length in order to be compared to each other because inDelphi can output the predictions of frequencies of the 1-bp insertions and deletions smaller than 60 bp. **c**. Indel frequencies of inDelphi predictions and NGS-measured experimental outcomes (SpCas9 only). Bar plot shows the frequencies of total deletions (red) and 1-bp insertions (blue) among overall on-target mutations. The colors of the sgRNA characters indicate how SpCas9 FE-induced 1-bp indels compared to SpCas9 in each sgRNA: purple indicates when SpCas9 dFE induced significantly more 1-bp indels than the SpCas9 control; orange indicates when SpCas9 iFE induced significantly more 1-bp indels than the SpCas9 control; mixed colors indicate that both SpCas9 iFE and SpCas9 dFE improved significantly over SpCas9; black indicates that there was no significant increase by SpCas9 dFE and SpCas9 iFE. +80 sgRNA indicates the sgRNA predicted by InDelphi for which more than 80% are deletion, and -80 sgRNA indicates the sgRNA for which less than 80% are deletion.

For SpCas9 dFE, we observed robust increases in 1-bp deletions using sgRNAs predicted by inDelphi to have predominant total deletions when used with SpCas9 (Figure 5c). Out of 20 sgRNA sequences tested, 11 were predicted to have >80% total deletion (+80 sgRNAs) by inDelphi and 9 were predicted to have <80% total deletion (−80 sgRNAs). When coupled with SpCas9 dFE, 9/11 of the +80 sgRNAs (82%) showed significantly increased 1-bp deletions experimentally, but only 2/9 of the -80 sgRNAs (22%) had significantly increased 1-bp deletions. This suggests that inDelphi predictions for sgRNA selection with SpCas9 editing could be repurposed for sgRNA selection for SpCas9 dFE.

However, the enhancement effect of 1-bp insertions for SpCas9 iFE did not correlate robustly with inDelphi predictions for specific sgRNAs (Figure 5c), despite the fact that SpCas9 iFE did increase 1-bp insertion rates in experiments using multiple sgRNAs compared to SpCas9. Thus, further work will be required to develop algorithms to design sgRNAs for SpCas9 iFE.

### Treatment of a pathogenic frameshift using frame editors

We used inDelphi to design a frame editing strategy to correct a pathogenic insertional mutation via a targeted deletion. A mutation in the *RAI1* gene causes Smith-Magenis syndrome^32–34^ (Figure 6a). To completely correct the RAI1^3103insC^ mutation by inducing a targeted 1-bp deletion, we selected RAI1-sgRNAs that were as close as possible to the RAI1^3103insC^ mutation site and for which dFE would likely to be effective (Figure 6b). InDelphi predicted that one particular RAI1-sgRNA would generate 92.2% total deletions (+80 sgRNAs) and based on the results above, we believed that this design has high potential to guide a precise 1-bp deletion at the targeted locus (Figure 6c).

**Figure 6.**
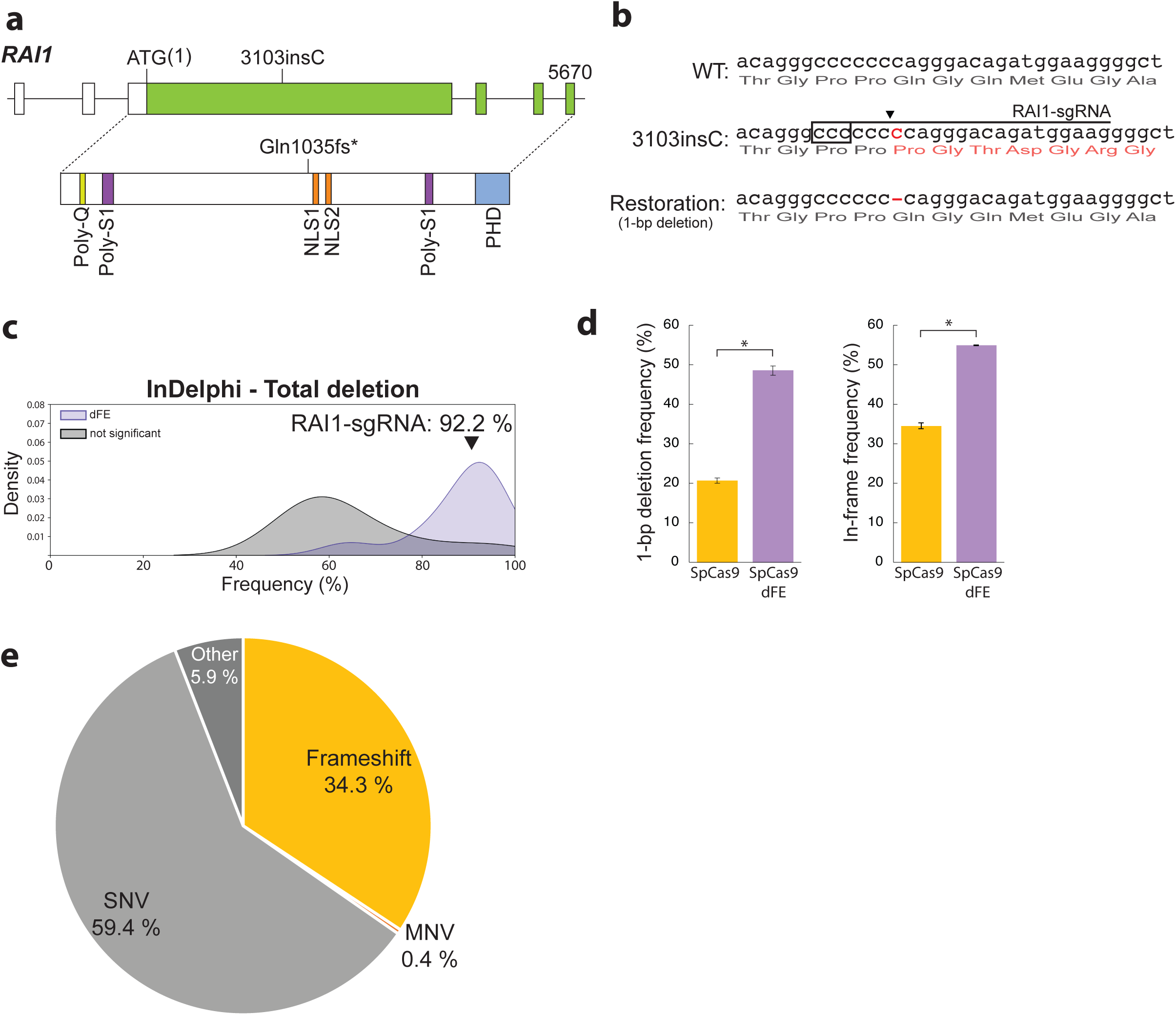
Correction of pathogenic mutation using FEs. Correction of pathogenic RAI1 mutation in HEK293T cells. **a**. Schematic of 3103insC mutation in RAI1 that causes Smith-Magenis syndrome^34^. **b**. RAI1^3103insC^ mutation and accurate correction by 1-bp deletion. The sgRNAs targeting RAI1^3103insC^ were designed around the mutated position, as indicated by the underline. Boxes indicate PAM sequence, black arrowhead indicates cleavage position, and red letter indicates inserted cytosine. **c**. Univariate distribution of total deletion predicted by inDelphi. The x-axis shows the predicted total deletion, and the y-axis shows the probability density calculated by kernel density estimation (KDE). Purple area indicate the sgRNAs having significant differences between the frequency of 1-bp deletions with SpCas9 dFE and SpCas9, and black area indicate the sgRNAs with no significant differences. Black arrowhead in the graph shows the frequency of total deletions of RAI1 sgRNA predicted by inDelphi. **d**. The RAI1^3103insC^ allele of HEK293T cells was restored using editing with SpCas9 and SpCas9 dFE. Shown are frequencies of therapeutic correction, which is 1-bp deletion and in-frame mutation to correct a frameshift disrupting proper protein translation, in on-target overall mutations. Data are expressed as mean□±□s.e.m. of n = 3 biological replicates. *P<0.05 (*Welch’s t-test*). **f**. Classification of pathogenic human gene variants by type in dbSNP (Total variants: 79083).

In RAI1^3103insC^ HEK293T cells, SpCas9 dFE with RAI1-sgRNA increased the frequency of 1-bp deletions by 2.3-fold, and increased the frequency of in-frame correction by 1.6-fold, compared with SpCas9 (Figure 6d). Thus, by using an sgRNA predicted by inDelphi to induce a high frequency of total deletions, we were able to increase the frequency of 1-bp deletion using SpCas9 dFE to correct the RAI1^3103insC^ mutation.

There are many examples of pathogenic frameshift mutations: the frequency of frameshift mutation entries in dbSNP is about 0.07%, but when these are filtered by those frameshift mutations which are pathogenic, this percentage rises to 34% of the total pathogenic entries (Figure 6e). Therefore, FEs may offer a new option for gene therapy against genetic diseases caused by frameshift mutations.

## Discussion

In this study, we developed FEs that shift the distribution of mutations caused by SpCas9 (and variants) towards 1-bp indels, resulting in precise 1-bp frameshifts. Since the frequency of overall mutations by FEs, including off-target effects, did not change significantly compared with those caused by SpCas9, the efficiency of genome editing with our FEs remain dictated by the cleavage activity of the Cas9 nuclease or its variants.

Comparisons between FEs constructed using SpCas9 and its variants revealed differences in the distribution of mutation types and the frequency of 1-bp indels (Figure S5). This diversity indicates that the DNA cleavage ends may be altered by the choice of sgRNA and Cas9 variant^21,22,35^, such as LZ3 Cas9. Thus, there should be opportunities to further optimize genome-editing outcomes with other Cas9 variants in the future. Many Cas9 variants have already been developed, such as PAM-less^36–38^ or high-fidelity Cas9^21,30,39–41^ versions. However, alternative-cleavage Cas9 variants should also be of interest for increasing the precision of CRISPR frame editing.

Algorithms for predicting Cas9-mediated mutations based on sgRNA sequences have been developed using data from large-scale sgRNA screening in various cell lines^12–15^. We used some of these algorithms to design sgRNAs that demonstrate enhanced frameshifting when used with our FEs. Future studies specifically focused on combining sgRNA libraries with FEs for frameshifting across diverse cell lines and target loci would enable FE-specific algorithms. Such algorithms, combined with further optimization of the enzyme variants used in FEs, could lead to enhanced optimization of editing outcomes.

One unanticipated finding in the present study was that T4pol could increase the frequency of 1-bp deletions and thereby reduce the number and types of other mutations. We hypothesized that T4pol induces 1-bp deletions by suppressing DSB-end resection; this is supported by a recent report^42^, which showed that fusing Cas9 with *Escherichia coli* DNA polymerase I and Klenow fragments increases the frequency of 1-bp deletions^42^, an observation that consistent with the results we obtained with dFE.

Frame editing is highly effective for gene disruption and frameshift correction because it ensures that the smallest predictable mutations can be introduced. The ±1-bp frameshifts introduced by FE can correct frameshifts either by causing the mutation to revert to the normal genotype or by making the mutations small and in-frame. Potential approaches for treating genetic disorders caused by frameshifts are already being evaluated in clinical trials; for example, exon skipping is one promising method for muscular dystrophy^43^. However, this technique can only be used for genes that do not lose their function when the exons are removed (e.g., structural proteins and repetitive regions). Our FEs, on the other hand, can expand the range of genetic diseases that could be treated.

Frame editing has applications beyond gene therapy. For example, FEs could improve CRISPR screens because they efficiently introduce the smallest frameshifts to result in knockouts. This characteristic could increase the resolution of tiling screening or reduce the number of sgRNAs for genome-wide screening. Another example is using accurate mutations by FEs to improve DNA writers. Self-targeting guide RNAs^44–46^, such as our previously developed mSCRIBE^45^, can record cellular phenomena *in vivo* over time, by inscribing changes in DNA sequences. Loveless *et al*. combined a self-targeting guide RNA with terminal deoxynucleotidyl transferase (TdT)^47^, which belongs to the X-family of DNA polymerases, to develop a DNA recorder that avoids the loss of recorded information by introducing insertions continuously. Combining FE with self-targeting sgRNA would allow the number of biological events to be determined based on the length of the sequence, since the recorded sequence would change by ∼1-bp each time a recording occurs.

This study provides a new method for introducing certain mutations by genome editing without a donor template. The FEs described here represent an important step in the development of genome editing by Cas9 as they point to a mechanism for controlling the length of the insertion or deletion. We anticipate that as research on frame editing technologies continues to advance, fully controllable gene modifications could be introduced without template.

## Methods

### Plasmid construction

Cas9 and FE expression vectors were constructed using the pX330A and pX330S vectors contained in Multiplex CRISPR/Cas9 Assembly System Kit (Kit #1000000055, Addgene)^48^ with some modifications. The DSB-end processing enzymes were either DNA synthesized with gBlocks (IDT) or reverse-transcribed from a natural source, and were inserted at the C-terminal ends of Cas9 in the pX330A and pX330S vectors. For Cas9 variants, the pX330A vector was modified by using a primer to introduce the R691A mutation. LZ3 Cas9 was obtained from Addgene (Plasmid #140561). Vectors expressing multiple sgRNAs were constructed using the Multiplex CRISPR/Cas9 Assembly System. Oligonucleotides for sgRNA templates were synthesized, annealed, and inserted into the modified pX330A and pX330S vectors, then cloned as an sgRNA cassette into pcDNA3.1(−) vector. Golden Gate assembly was used to assemble the constructed vectors into the all-in-one CRISPR/Cas9 vectors. The RAI1 sgRNA expressing vectors were constructed with pcDNA3.1(−) vector, which inserted sgRNA cloning cassette. The sequences of FEs are shown in Supplementary Sequences 1.

### sgRNA design

The 7 sgRNAs with various distributions of mutations were chosen based on the study of mutation profiling by genome editing^49,50^. The other 13 sgRNAs were selected from the 64,751 GeCKO library^51^ to assess the ability of frame editing for sgRNAs inducing various mutations. The sgRNAs targeting a gene ranked within the top 1,000 essential genes in any cell type (Supplementary Table 2) were discarded from the list of candidates to avoid bias resulting from the lethality of mutating those genes. inDelphi^14^ (scikit-learn v0.20.0 models, for HEK293) was then applied to select only the gRNA that would show definite distributions on indels. The sgRNA targets whose “Precision” value was higher than 0.35 were selected as the final candidates. The more detailed explanations and Python script for the selection process were uploaded to our GitHub repository (https://github.com/KazukiNakamae/Frame_Editor_sgRNA_selection).

### Cell culture

HEK293T cells obtained from ATCC were maintained in Dulbecco’s modified Eagle’s medium (DMEM) GlutaMAX (Gibco) supplemented with 10% (v/v) fetal bovine serum (Corning). The cells were cultured at 37 °C with 5% CO_2_. The cell line was tested as being negative for mycoplasma contamination using the LookOut Mycoplasma PCR Detection Kit (Sigma-Aldrich) and Jumpstart™ Taq DNA Polymerase (Sigma-Aldrich).

### Transfection for frame editing

The plasmids were transfected using FuGENE HD (Promega) and Opti-MEM (Gibco), then genomic DNA was collected after 48 hours using DNAzol™ Reagent (Invitrogen), according to the manufacturer’s instructions. Equal amounts of the plasmids were mixed in experiments for collecting the RAI1^3103insC^ mutation (Cas9 or dFE/sgRNA vector) and in experiments for genome editing by FE (Cas9 or FE/GFP/sgRNA vector). In the experiments shown in Fig. 1, S1a, S5, and S7, 3 × 10^4^ HEK293T cells were transfected with 200 ng of plasmids using a 96-well plate (i.e., without sorting). In all other experiments, 3 × 10^5^ HEK293T cells were transfected with 2000 ng of plasmids using a 12-well plate, then GFP-fluorescent cells were sorted by BD FACSAria™ IIIu (BD Biosciences), and the genomic DNA was immediately extracted from the collected cells.

### TIDE analysis

Genomic PCR was performed using NEBNext High-Fidelity 2X PCR Master Mix (New England Biolabs) with the primers listed in Supplementary Table 1. The PCR products were run on a 1.5% (wt/vol) agarose gel. The target band was cut out and extracted by QIAquick Gel Extraction Kit (Qiagen). The purified PCR products were mixed with primers for sequencing in order to submit to Sanger sequencing service provided by GENEWIZ/Azenta Life Sciences (https://www.genewiz.com/). Sanger sequencing data downloaded in ab1 format were analyzed by TIDE^52^ (http://shinyapps.datacurators.nl/tide/). Data are shown in Supplementary Table 2. The mutation frequencies with a P-value threshold of 0.001 or higher were excluded.

### NGS and data analysis

Genomic PCR was performed using KAPA HiFi HotStart DNA Polymerase (Roche) with the primers listed in Supplementary Table 1. The PCR products were run on a 1.5% (wt/vol) agarose gel. The target band was cut out and extracted with the QIAquick Gel Extraction Kit (Qiagen). The purified PCR products were mixed with multiple samples containing different sequences, then concentrated using isopropanol precipitation with GlycoBlue Coprecipitant (Invitrogen) by submitting to Amplicon-EZ, which is the service for next-generation sequencing (NGS), provided by GENEWIZ/Azenta Life Sciences (https://www.genewiz.com/). The NGS data were analyzed using CRISPResso2^53^ (https://github.com/pinellolab/CRISPResso2). Running mode was set to CRISPRessoPooled, and quantification window size was set to 2. Of the output by CRISPResso2, the frequency of inserted bases in 1-bp insertion and microhomology-associated deletion was analyzed using Alleles_frequency_table_around_sgRNA_*, and the on-target and off-target mutations in Figs. 4d and 4e were used. Frequencies of mutations were calculated as described in SAMPLES_QUANTIFICATION_SUMMARY_*, while all other analyses were done using Indel_histogram. These mutation frequencies were organized in Microsoft Excel and Python, and then visualized. Data are shown in Supplementary Table 3. The Symmetrized KL divergence in Fig. S3 was calculated and visualized using the homemade script (https://github.com/KazukiNakamae/Frame_Editor_KLD_calculation).

### Off-target site selection

Potential off-target editing sites were searched using CRISPOR (http://crispor.tefor.net/). We selected off-target sites according to their MIT scores. Off-target sites were calculated for 20 sgRNAs using Fig 2, 3, 4 and 5, and the 9 sites having the highest MIT off-target scores (Supplementary Table 2).

### Generation of HEK293T cells containing RAI1^3103insC^ mutation

HEK293T cells with the RAI1^3103insC^ mutation were generated by introducing the plasmid expressing Cas9 and sgRNA with single-stranded oligodeoxynucleotides (Supplementary Sequence 2) by FuGENE HD (Promega). The Cas9-expression plasmid with the sgRNA sequence for generating the RAI1^3103insC^ mutation and the GFP/puromycin expression plasmid (70 ng of each plasmid) were mixed with 0.3 μL of single-strand oligo nucleotide at 10 μM. Twenty-four hours after lipofection, the medium was replaced with DMEM with 10 ng/μL puromycin and treated for 2 days. The drug-selected cells were cloned by limiting dilution in a 96-well plate, then a clone containing the RAI1^3103insC^ mutation in all detective alleles was identified by Sanger sequencing.

### Statistical analysis

Statistical analysis was performed using Microsoft Excel, R script, and Python. All data are presented as mean ± s.e.m. All measurements were taken from biological replicates. Statistical significance was determined by *Welch’s t-test*.

## Supporting information

Supplementary_sequences

## Acknowledgments

We would like to acknowledge the technical assistance of the Koch Institute Flow Cytometry Core for cell sorting, the general assistance of Karen Pepper (Massachusetts Institute of Technology) for editing the manuscript, and Ky Lowenhaupt (Massachusetts Institute of Technology) for overall management. We also thank Feng Zhang (Broad Institute) for the provision of reagents through Addgene. S.N. would like to thank the late Sheila Hoffman and her family for their help and support during his time in United States. S.N. acknowledge the Overseas Research Fellowship at the Japan Society for the Promotion of Science and Postdoctoral Fellowship at Uehara Memorial Foundation. This work was also supported by a grant from NIH (T.K.L U01CA250554), and partially by Cancer Center Support (core) Grant P30-CA14051 from the NCI. S.N. and T.K.L. filed a patent application for FEs^54^.

## Figure Legends

**Figure S1.**
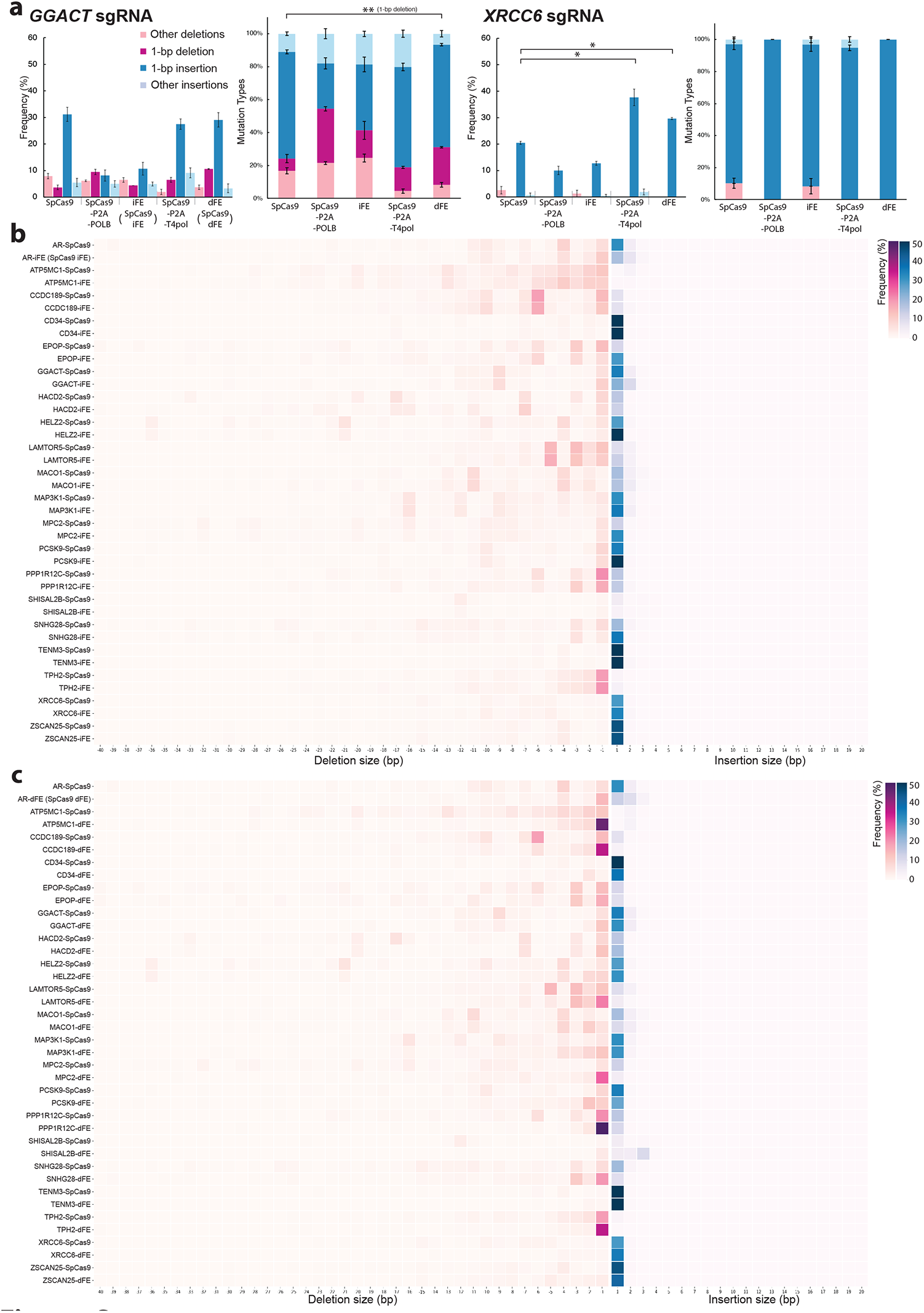
Editing outcomes obtained with SpCas9, SpCas9 iFE, and SpCas9 dFE. **a**. The indel efficiency in HEK293T cells was measured by TIDE analysis. Shown for each sgRNA are the frequencies of mutation types in sequence traces (left), and the frequencies of mutation types among on-target overall mutations (right). TIDE analysis was calculated by setting the indel size range to 10 and excluding those with P ≥ 0.001. Mean ± s.e.m. of n = 3 independent biological replicates. 1-bp insertion: *P<0.05, 1-bp deletion: **P<0.05 (*Welch’s t-test*). **b-c**. Comparison of mutations in HEK293T cells induced by SpCas9 and SpCas9 iFE **(b)** and by SpCas9 and SpCas9 dFE **(c)**. Shown are the distributions of indel size. The heatmap indicates frequencies of insertions (blue) and deletions (red) in total reads. For each sgRNA, the y-axis is SpCas9 (top) and SpCas9 iFE (bottom), and the x-axis is indel size. Target genes of sgRNA are shown in the upper left. Only transfected cells were collected by GFP+ sorting. Data are expressed as means□±□s.e.m. of n = 3 biological replicates.

**Figure S2.**
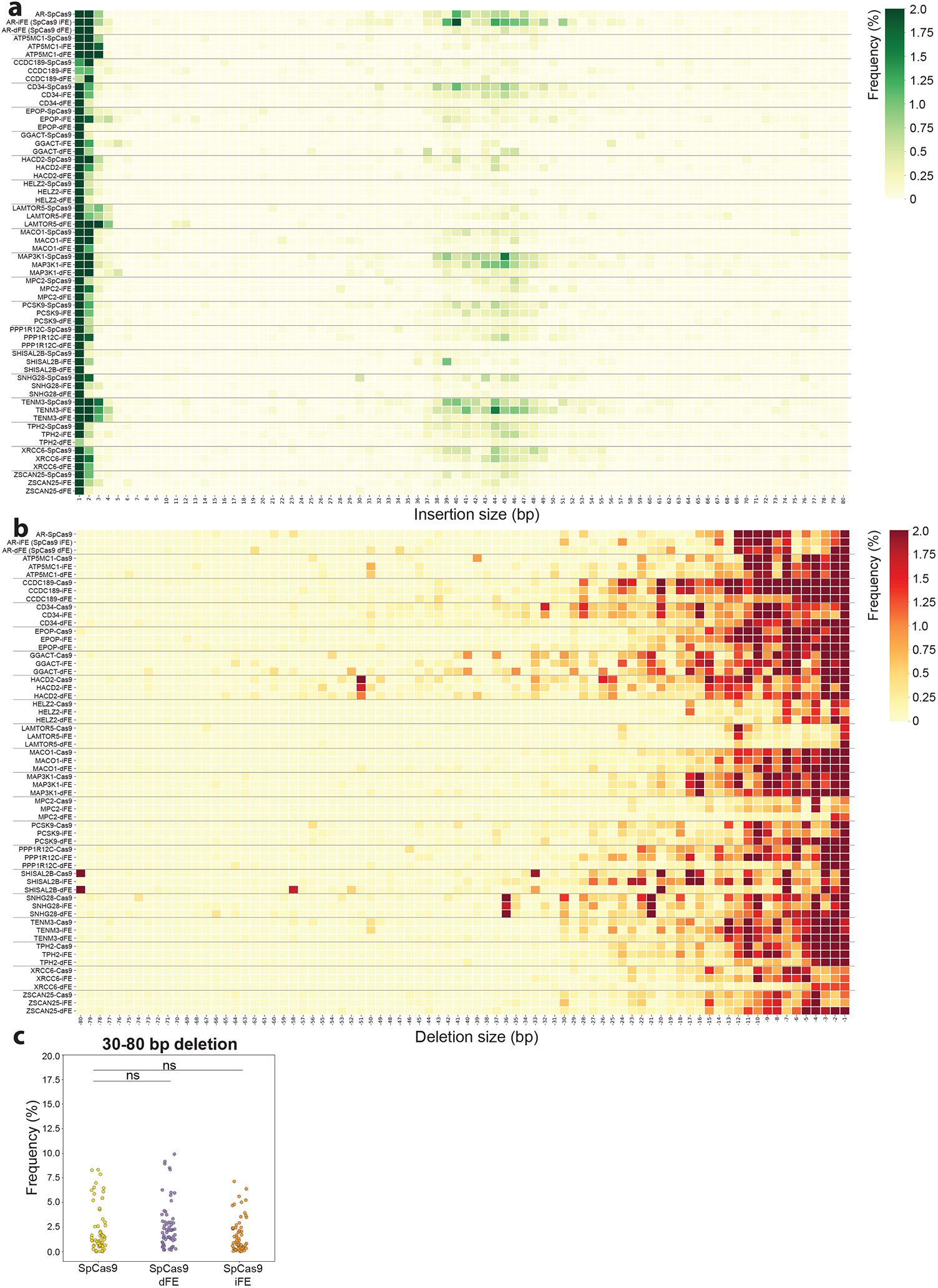
Large-size mutation outcomes for all sgRNAs using SpCas9 iFE and SpCas9 dFE. **a-b**. Comparison of large size mutations induced by SpCas9, SpCas9 iFE, and SpCas9 dFE in HEK293T cells. Shown are the distributions of indel size. The heatmap indicates the frequency of insertions (blue, **a**) and deletions (red, **b**) in total reads. Shown for each sgRNA: the y-axis is SpCas9 (top), SpCas9 iFE (middle), and SpCas9 dFE (bottom); and the x-axis is indel size. **c**. Comparison of large deletions induced by SpCas9, SpCas9 iFE, and SpCas9 dFE in HEK293T cells. Total frequency of 30−80-bp indels for each sgRNA. Only transfected cells were collected by GFP+ sorting. Data are expressed as means□±□s.e.m. of n = 3 biological replicates. *P<0.05 (*Welch’s t-test*).

**Figure S3.**
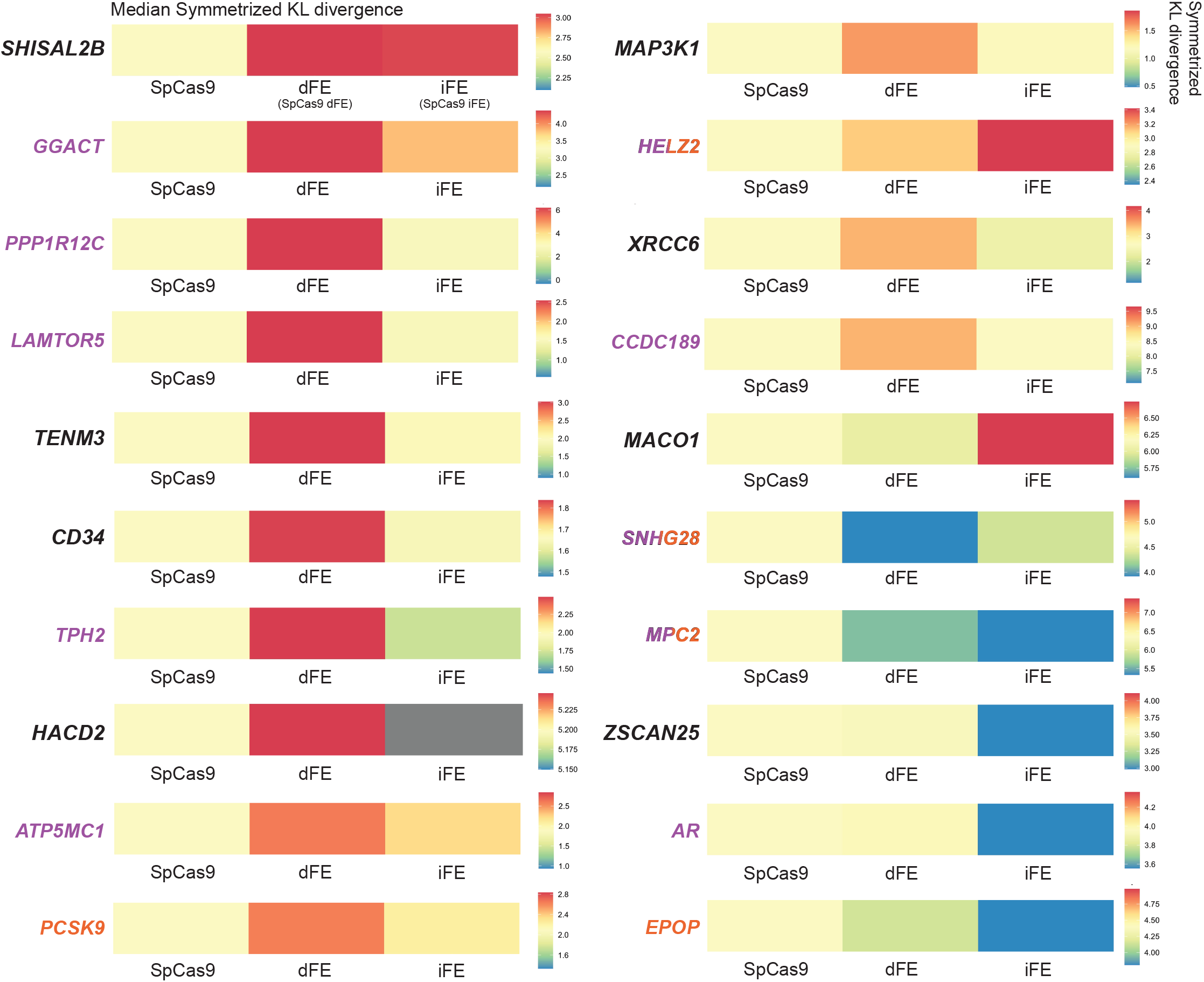
Comparison of inDelphi predictions and FE outcomes by KL divergence. Symmetrized KL divergence to visualize the difference between distributions of mutations introduced by SpCas9, SpCas9 dFE, and SpCas9 iFE and mutations predicted by inDelphi. The KL divergence indicates the median value of each replicate. The colors of the sgRNA characters indicate how SpCas9 FE-induced 1-bp indels compared to SpCas9 in each sgRNA: purple indicates when SpCas9 dFE induced significantly more 1-bp indels than the SpCas9 control; orange indicates when SpCas9 iFE induced significantly more 1-bp indels than the SpCas9 control; mixed colors indicate that both SpCas9 iFE and SpCas9 dFE improved significantly over SpCas9; black indicates that there was no significant increase by SpCas9 dFE and SpCas9 iFE.

**Figure S4.**
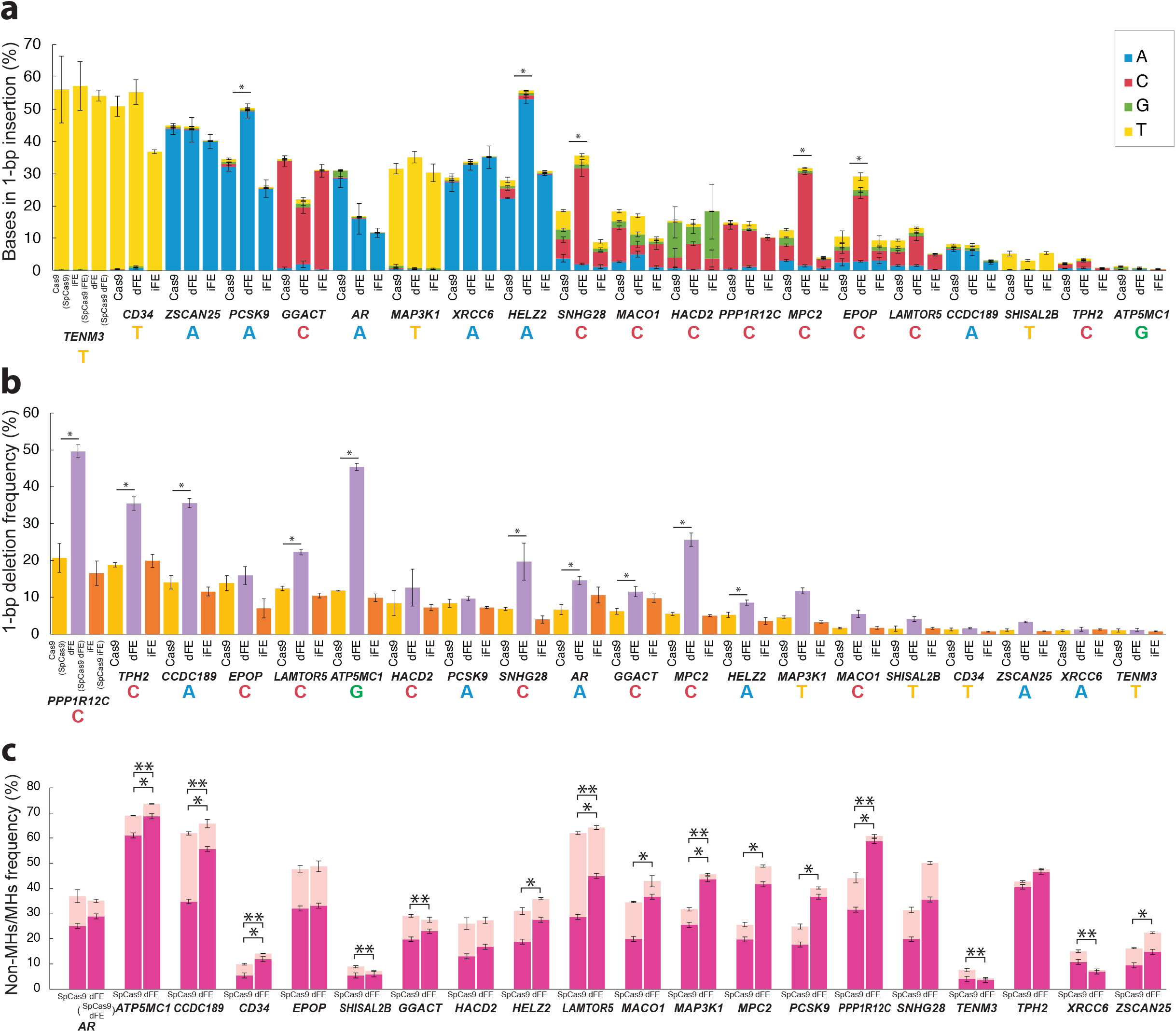
Analysis of the mutation types depending on the target sequence. **a-b**. Frequencies of inserted bases **(a)** and 1-bp deletion **(b)** for all sgRNAs. The bases on staggered ends are shown at the bottom of the x-axis. **c**. Frequency of non-MHs (microhomologies) / MHs. The mutation reads with deletions of 15-bp or less in length were analyzed, and ≥2-bp homology was classified as MHs. Only transfected cells were collected by GFP+ sorting. Data are expressed as mean□±□s.e.m. of n = 3 biological replicates. *P<0.05 (*Welch’s t-test*).

**Figure S5.**
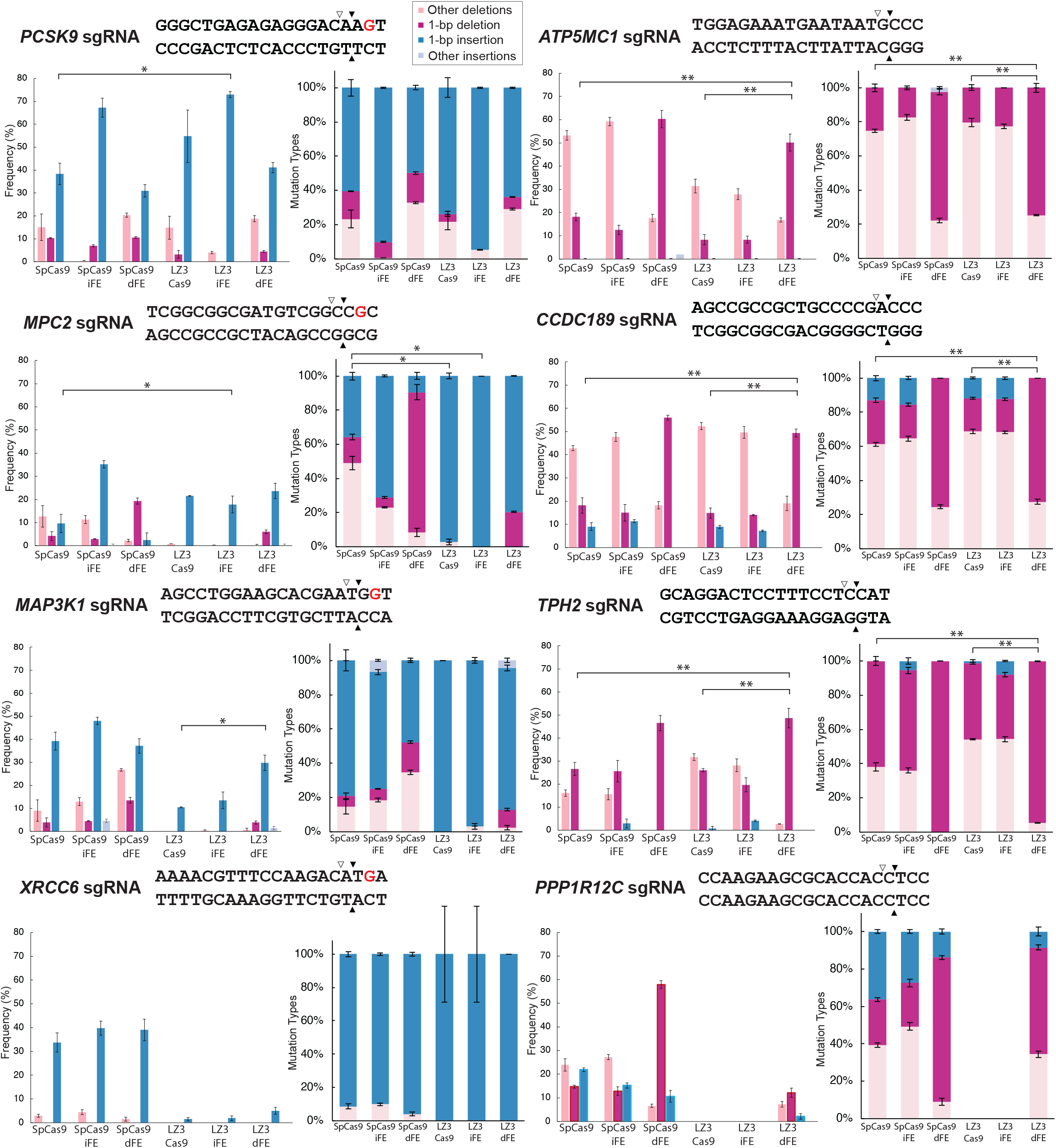
Effect of LZ3 FEs. Indel efficiency of LZ3 Cas9 and LZ3 FEs in HEK293T cells was measured by TIDE analysis. Shown for each sgRNA are the frequencies of mutation types in sequence traces (left) and the frequencies of mutation types among on-target overall mutations (right). Black triangles indicate blunt end cut sites, and white triangles indicate staggered end cut sites. Red letters indicate a guanine in the -2 PAM that increases the frequency of 1-bp insertions with LZ3 Cas9. TIDE analysis was calculated by setting the indel size range 1 to 10 and excluding those with P ≥ 0.001. Mean ± s.e.m. of n = 3 independent biological replicates. 1-bp insertion: *P<0.05, 1-bp deletion: **P<0.05 (*Welch’s t-test*).

**Figure S6.**
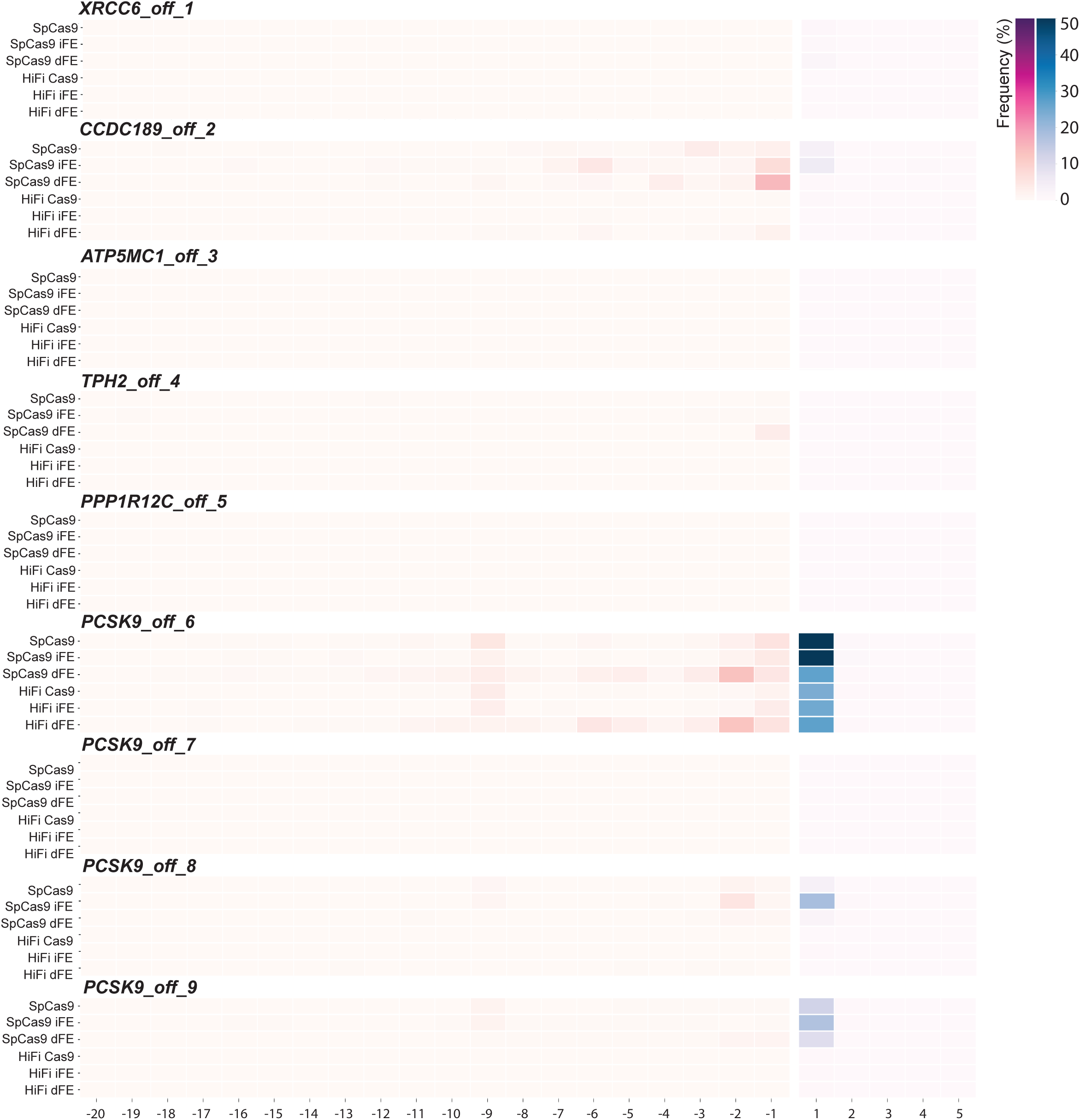
Distribution of off-target sizes induced by SpCas9 FE and HiFi FE. Comparison of distribution of off-target mutations induced by SpCas9, SpCas9 iFE, SpCas9 dFE, HiFi Cas9, HiFi iFE, and HiFi dFE in HEK293T cells. Target genes of sgRNA are shown in the upper left. The heatmap indicates frequencies of insertions (blue) and deletions (red) in total reads. For each sgRNA, the y-axis indicates Cas9 and FEs, and the x-axis corresponds to indel size. Only transfected cells were collected by GFP+ sorting. Data are expressed as mean□±□s.e.m. of n = 3 biological replicates. *P<0.05 (*Welch’s t-test*).

**Figure S7.**
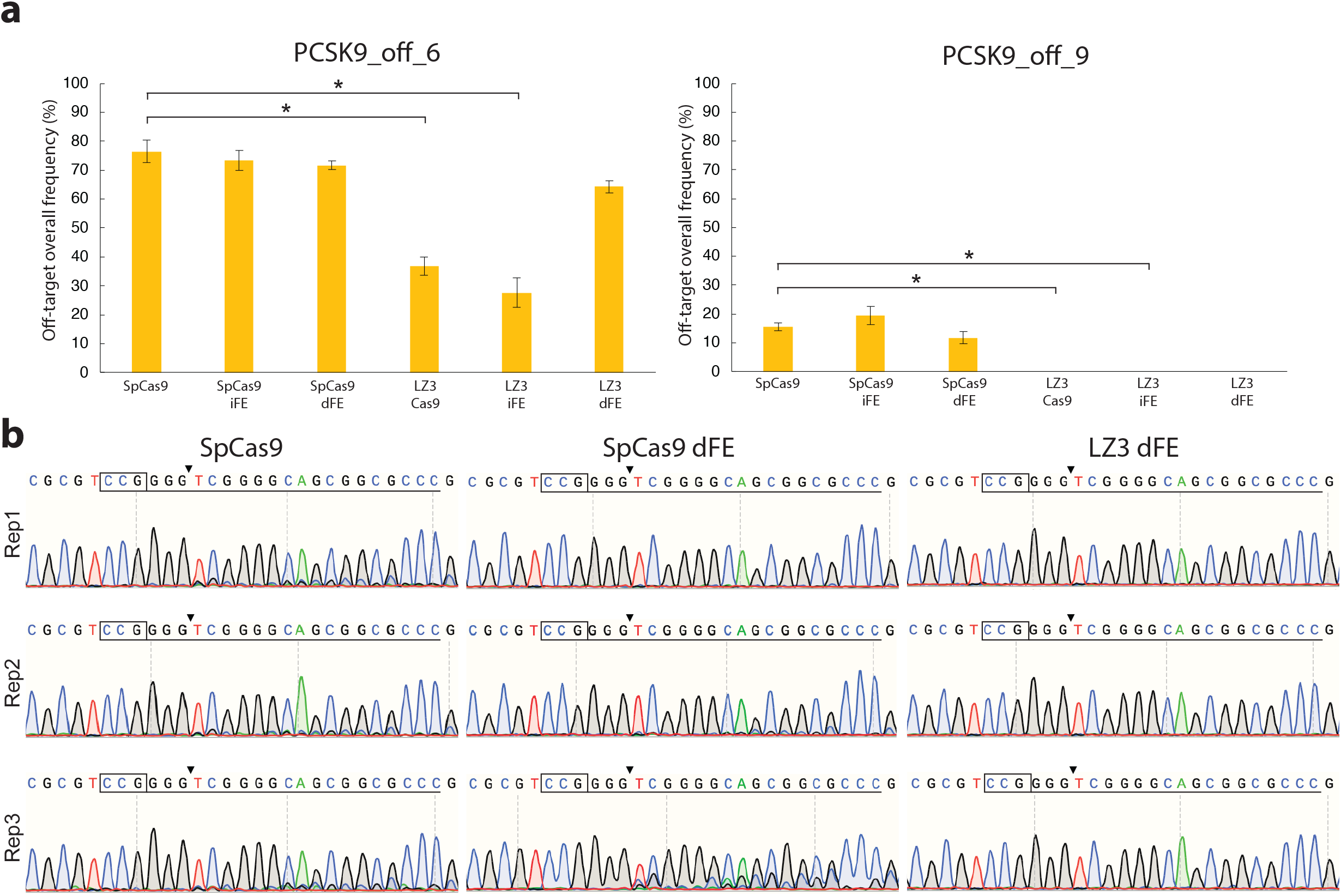
Off-target effect with LZ3 Cas9 and LZ3 FEs. **a**. Off-target editing of predicted off-target sites PCSK9_off_6 and PCSK9_off_9 for SpCas9, SpCas9 FEs, LZ3 Cas9, and LZ3 FEs. **b**. Off-target editing of the CCDC189_off_2 for SpCas9, SpCas9 dFE, and LZ3 dFE. Since there were multiple identical sequences in a short region, analysis by TIDE was not possible. Thus, the sequence data is shown. The y-axis shows the biological replicates, and the x-axis shows the editing tools used (SpCas9, SpCas9 dFE, and LZ3 dFE).

