## Supplementary_sequences for "Frame Editors for Precise, Template-Free Frameshifting"

**Supplementary Sequences 1.**

Coding sequences for frame editing

SpCas9-P2A-POLB:

atggactataaggaccacgacggagactacaaggatcatgatattgattacaaagacgatgacgataagatggccccaaagaagaagcggaaggtcggtatccacggagtcccagcagccgacaagaagtacagcatcggcctggacatcggcaccaactctgtgggctgggccgtgatcaccgacgagtacaaggtgcccagcaagaaattcaaggtgctgggcaacaccgaccggcacagcatcaagaagaacctgatcggagccctgctgttcgacagcggcgaaacagccgaggccacccggctgaagagaaccgccagaagaagatacaccagacggaagaaccggatctgctatctgcaagagatcttcagcaacgagatggccaaggtggacgacagcttcttccacagactggaagagtccttcctggtggaagaggataagaagcacgagcggcaccccatcttcggcaacatcgtggacgaggtggcctaccacgagaagtaccccaccatctaccacctgagaaagaaactggtggacagcaccgacaaggccgacctgcggctgatctatctggccctggcccacatgatcaagttccggggccacttcctgatcgagggcgacctgaaccccgacaacagcgacgttgacaagctgttcatccagctggtgcagacctacaaccagctgttcgaggaaaaccccatcaacgccagcggcgtggacgccaaggccatcctgtctgccagactgagcaagagcagacggctggaaaatctgatcgcccagctgcccggcgagaagaagaatggcctgttcggaaacctgattgccctgagcctgggcctgacccccaacttcaagagcaacttcgacctggccgaggatgccaaactgcagctgagcaaggacacctacgacgacgacctggacaacctgctggcccagatcggcgaccagtacgccgacctgtttctggccgccaagaacctgtccgacgccatcctgctgagcgacatcctgagagtgaacaccgagatcaccaaggcccccctgagcgcctctatgatcaagagatacgacgagcaccaccaggacctgaccctgctgaaagctctcgtgcggcagcagctgcctgagaagtacaaagagattttcttcgaccagagcaagaacggctacgccggctacattgacggcggagccagccaggaagagttctacaagttcatcaagcccatcctggaaaagatggacggcaccgaggaactgctcgtgaagctgaacagagaggacctgctgcggaagcagcggaccttcgacaacggcagcatcccccaccagatccacctgggagagctgcacgccattctgcggcggcaggaagatttttacccattcctgaaggacaaccgggaaaagatcgagaagatcctgaccttccgcatcccctactacgtgggccctctggccaggggaaacagcagattcgcctggatgaccagaaagagcgaggaaaccatcaccccctggaacttcgaggaagtggtggacaagggcgcttccgcccagagcttcatcgagcggatgaccaacttcgataagaacctgcccaacgagaaggtgctgcccaagcacagcctgctgtacgagtacttcaccgtgtataacgagctgaccaaagtgaaatacgtgaccgagggaatgagaaagcccgccttcctgagcggcgagcagaaaaaggccatcgtggacctgctgttcaagaccaaccggaaagtgaccgtgaagcagctgaaagaggactacttcaagaaaatcgagtgcttcgactccgtggaaatctccggcgtggaagatcggttcaacgcctccctgggcacataccacgatctgctgaaaattatcaaggacaaggacttcctggacaatgaggaaaacgaggacattctggaagatatcgtgctgaccctgacactgtttgaggacagagagatgatcgaggaacggctgaaaacctatgcccacctgttcgacgacaaagtgatgaagcagctgaagcggcggagatacaccggctggggcaggctgagccggaagctgatcaacggcatccgggacaagcagtccggcaagacaatcctggatttcctgaagtccgacggcttcgccaacagaaacttcatgcagctgatccacgacgacagcctgacctttaaagaggacatccagaaagcccaggtgtccggccagggcgatagcctgcacgagcacattgccaatctggccggcagccccgccattaagaagggcatcctgcagacagtgaaggtggtggacgagctcgtgaaagtgatgggccggcacaagcccgagaacatcgtgatcgaaatggccagagagaaccagaccacccagaagggacagaagaacagccgcgagagaatgaagcggatcgaagagggcatcaaagagctgggcagccagatcctgaaagaacaccccgtggaaaacacccagctgcagaacgagaagctgtacctgtactacctgcagaatgggcgggatatgtacgtggaccaggaactggacatcaaccggctgtccgactacgatgtggaccatatcgtgcctcagagctttctgaaggacgactccatcgacaacaaggtgctgaccagaagcgacaagaaccggggcaagagcgacaacgtgccctccgaagaggtcgtgaagaagatgaagaactactggcggcagctgctgaacgccaagctgattacccagagaaagttcgacaatctgaccaaggccgagagaggcggcctgagcgaactggataaggccggcttcatcaagagacagctggtggaaacccggcagatcacaaagcacgtggcacagatcctggactcccggatgaacactaagtacgacgagaatgacaagctgatccgggaagtgaaagtgatcaccctgaagtccaagctggtgtccgatttccggaaggatttccagttttacaaagtgcgcgagatcaacaactaccaccacgcccacgacgcctacctgaacgccgtcgtgggaaccgccctgatcaaaaagtaccctaagctggaaagcgagttcgtgtacggcgactacaaggtgtacgacgtgcggaagatgatcgccaagagcgagcaggaaatcggcaaggctaccgccaagtacttcttctacagcaacatcatgaactttttcaagaccgagattaccctggccaacggcgagatccggaagcggcctctgatcgagacaaacggcgaaaccggggagatcgtgtgggataagggccgggattttgccaccgtgcggaaagtgctgagcatgccccaagtgaatatcgtgaaaaagaccgaggtgcagacaggcggcttcagcaaagagtctatcctgcccaagaggaacagcgataagctgatcgccagaaagaaggactgggaccctaagaagtacggcggcttcgacagccccaccgtggcctattctgtgctggtggtggccaaagtggaaaagggcaagtccaagaaactgaagagtgtgaaagagctgctggggatcaccatcatggaaagaagcagcttcgagaagaatcccatcgactttctggaagccaagggctacaaagaagtgaaaaaggacctgatcatcaagctgcctaagtactccctgttcgagctggaaaacggccggaagagaatgctggcctctgccggcgaactgcagaagggaaacgaactggccctgccctccaaatatgtgaacttcctgtacctggccagccactatgagaagctgaagggctcccccgaggataatgagcagaaacagctgtttgtggaacagcacaagcactacctggacgagatcatcgagcagatcagcgagttctccaagagagtgatcctggccgacgctaatctggacaaagtgctgtccgcctacaacaagcaccgggataagcccatcagagagcaggccgagaatatcatccacctgtttaccctgaccaatctgggagcccctgccgccttcaagtactttgacaccaccatcgaccggaagaggtacaccagcaccaaagaggtgctggacgccaccctgatccaccagagcatcaccggcctgtacgagacacggatcgacctgtctcagctgggaggcgacaaaaggccggcggccacgaaaaaggccggccaggcaaaaaagaaaaagataagcgctggaagcggagctactaacttcagcctgctgaagcaggctggagacgtggaggagaaccctggaccttctaagcgaaaggctccgcaggagacccttaacggtggcatcacggatatgctgacggagctggctaattttgaaaaaaatgtttcacaagctatccacaagtacaacgcctacagaaaagctgcttctgtaatagccaaatatcctcataaaatcaaatcaggggccgaggccaaaaaactgccgggggtggggacgaaaatcgcagaaaaaatagatgagtttttggcgacgggtaaattgcgaaagctggaaaagatcaggcaggacgacacctcttctagtattaatttcctcacgagagtaagcggtatcggacctagtgccgccaggaagttcgtagatgaggggattaagacattggaagatctccggaagaacgaggacaagttgaatcatcaccagcgaataggacttaaatatttcggagacttcgaaaagcgcattccacgagaggagatgctgcaaatgcaagacatagtgcttaatgaagtcaaaaaagtagacagcgagtacatagctactgtctgcggatcttttaggcggggtgctgagtcttctggtgatatggatgtactgctgactcatccatcttttacaagcgagagcactaagcaaccaaaactgcttcaccaagtcgtcgagcaactgcagaaggtacactttatcactgacactctgtctaaaggtgaaacgaagttcatgggcgtctgccagttgccgagtaaaaatgatgaaaaagaatatccgcatcggagaatagacatccgactcatcccgaaggaccaatactactgtggggtcttgtacttcacgggctcagacattttcaacaagaatatgagggcacacgccctggaaaagggttttacaataaatgagtacacgatcagacctctgggagtcaccggggttgcgggagagccgcttccggttgactcagaaaaggacattttcgactatatccagtggaaatacagagaaccaaaggacagaagtgagggtggttctccaaagaagaagcggaaggtctag

3xFlagtag

Cas9

40SVNLS

Nucleoplasmin NLS

P2A linker

Polymerase beta

SpCas9-P2A-T4pol:

atggactataaggaccacgacggagactacaaggatcatgatattgattacaaagacgatgacgataagatggccccaaagaagaagcggaaggtcggtatccacggagtcccagcagccgacaagaagtacagcatcggcctggacatcggcaccaactctgtgggctgggccgtgatcaccgacgagtacaaggtgcccagcaagaaattcaaggtgctgggcaacaccgaccggcacagcatcaagaagaacctgatcggagccctgctgttcgacagcggcgaaacagccgaggccacccggctgaagagaaccgccagaagaagatacaccagacggaagaaccggatctgctatctgcaagagatcttcagcaacgagatggccaaggtggacgacagcttcttccacagactggaagagtccttcctggtggaagaggataagaagcacgagcggcaccccatcttcggcaacatcgtggacgaggtggcctaccacgagaagtaccccaccatctaccacctgagaaagaaactggtggacagcaccgacaaggccgacctgcggctgatctatctggccctggcccacatgatcaagttccggggccacttcctgatcgagggcgacctgaaccccgacaacagcgacgttgacaagctgttcatccagctggtgcagacctacaaccagctgttcgaggaaaaccccatcaacgccagcggcgtggacgccaaggccatcctgtctgccagactgagcaagagcagacggctggaaaatctgatcgcccagctgcccggcgagaagaagaatggcctgttcggaaacctgattgccctgagcctgggcctgacccccaacttcaagagcaacttcgacctggccgaggatgccaaactgcagctgagcaaggacacctacgacgacgacctggacaacctgctggcccagatcggcgaccagtacgccgacctgtttctggccgccaagaacctgtccgacgccatcctgctgagcgacatcctgagagtgaacaccgagatcaccaaggcccccctgagcgcctctatgatcaagagatacgacgagcaccaccaggacctgaccctgctgaaagctctcgtgcggcagcagctgcctgagaagtacaaagagattttcttcgaccagagcaagaacggctacgccggctacattgacggcggagccagccaggaagagttctacaagttcatcaagcccatcctggaaaagatggacggcaccgaggaactgctcgtgaagctgaacagagaggacctgctgcggaagcagcggaccttcgacaacggcagcatcccccaccagatccacctgggagagctgcacgccattctgcggcggcaggaagatttttacccattcctgaaggacaaccgggaaaagatcgagaagatcctgaccttccgcatcccctactacgtgggccctctggccaggggaaacagcagattcgcctggatgaccagaaagagcgaggaaaccatcaccccctggaacttcgaggaagtggtggacaagggcgcttccgcccagagcttcatcgagcggatgaccaacttcgataagaacctgcccaacgagaaggtgctgcccaagcacagcctgctgtacgagtacttcaccgtgtataacgagctgaccaaagtgaaatacgtgaccgagggaatgagaaagcccgccttcctgagcggcgagcagaaaaaggccatcgtggacctgctgttcaagaccaaccggaaagtgaccgtgaagcagctgaaagaggactacttcaagaaaatcgagtgcttcgactccgtggaaatctccggcgtggaagatcggttcaacgcctccctgggcacataccacgatctgctgaaaattatcaaggacaaggacttcctggacaatgaggaaaacgaggacattctggaagatatcgtgctgaccctgacactgtttgaggacagagagatgatcgaggaacggctgaaaacctatgcccacctgttcgacgacaaagtgatgaagcagctgaagcggcggagatacaccggctggggcaggctgagccggaagctgatcaacggcatccgggacaagcagtccggcaagacaatcctggatttcctgaagtccgacggcttcgccaacagaaacttcatgcagctgatccacgacgacagcctgacctttaaagaggacatccagaaagcccaggtgtccggccagggcgatagcctgcacgagcacattgccaatctggccggcagccccgccattaagaagggcatcctgcagacagtgaaggtggtggacgagctcgtgaaagtgatgggccggcacaagcccgagaacatcgtgatcgaaatggccagagagaaccagaccacccagaagggacagaagaacagccgcgagagaatgaagcggatcgaagagggcatcaaagagctgggcagccagatcctgaaagaacaccccgtggaaaacacccagctgcagaacgagaagctgtacctgtactacctgcagaatgggcgggatatgtacgtggaccaggaactggacatcaaccggctgtccgactacgatgtggaccatatcgtgcctcagagctttctgaaggacgactccatcgacaacaaggtgctgaccagaagcgacaagaaccggggcaagagcgacaacgtgccctccgaagaggtcgtgaagaagatgaagaactactggcggcagctgctgaacgccaagctgattacccagagaaagttcgacaatctgaccaaggccgagagaggcggcctgagcgaactggataaggccggcttcatcaagagacagctggtggaaacccggcagatcacaaagcacgtggcacagatcctggactcccggatgaacactaagtacgacgagaatgacaagctgatccgggaagtgaaagtgatcaccctgaagtccaagctggtgtccgatttccggaaggatttccagttttacaaagtgcgcgagatcaacaactaccaccacgcccacgacgcctacctgaacgccgtcgtgggaaccgccctgatcaaaaagtaccctaagctggaaagcgagttcgtgtacggcgactacaaggtgtacgacgtgcggaagatgatcgccaagagcgagcaggaaatcggcaaggctaccgccaagtacttcttctacagcaacatcatgaactttttcaagaccgagattaccctggccaacggcgagatccggaagcggcctctgatcgagacaaacggcgaaaccggggagatcgtgtgggataagggccgggattttgccaccgtgcggaaagtgctgagcatgccccaagtgaatatcgtgaaaaagaccgaggtgcagacaggcggcttcagcaaagagtctatcctgcccaagaggaacagcgataagctgatcgccagaaagaaggactgggaccctaagaagtacggcggcttcgacagccccaccgtggcctattctgtgctggtggtggccaaagtggaaaagggcaagtccaagaaactgaagagtgtgaaagagctgctggggatcaccatcatggaaagaagcagcttcgagaagaatcccatcgactttctggaagccaagggctacaaagaagtgaaaaaggacctgatcatcaagctgcctaagtactccctgttcgagctggaaaacggccggaagagaatgctggcctctgccggcgaactgcagaagggaaacgaactggccctgccctccaaatatgtgaacttcctgtacctggccagccactatgagaagctgaagggctcccccgaggataatgagcagaaacagctgtttgtggaacagcacaagcactacctggacgagatcatcgagcagatcagcgagttctccaagagagtgatcctggccgacgctaatctggacaaagtgctgtccgcctacaacaagcaccgggataagcccatcagagagcaggccgagaatatcatccacctgtttaccctgaccaatctgggagcccctgccgccttcaagtactttgacaccaccatcgaccggaagaggtacaccagcaccaaagaggtgctggacgccaccctgatccaccagagcatcaccggcctgtacgagacacggatcgacctgtctcagctgggaggcgacaaaaggccggcggccacgaaaaaggccggccaggcaaaaaagaaaaagataagcgctggaagcggagctactaacttcagcctgctgaagcaggctggagacgtggaggagaaccctggacctaaggaattctatattagtatcgagacagtgggaaacaacattgtagagcggtatatcgatgaaaacggcaaagaaaggactcgagaggtcgaataccttccgaccatgtttcgccactgtaaagaagaatctaagtacaaagatatttacgggaagaattgcgctccccaaaaatttccctccatgaaggatgctcgagactggatgaagcgcatggaggacataggtctcgaagcattggggatgaacgattttaagttggcttacatctccgacacttacgggtcagaaatagtgtacgataggaagttcgttcgcgtggcaaattgcgatatagaggtcactggagataagttcccggacccgatgaaggcggagtacgaaattgacgctataacacactatgactcaatcgacgaccggttctatgtatttgacctgctcaattccatgtacgggtctgtaagcaagtgggacgctaaactcgcggctaaacttgactgtgaaggaggggatgaagtacctcaggaaatcttggacagggtaatctacatgccctttgacaatgaacgagatatgcttatggagtacattaatttgtgggagcaaaagcgccccgcaatatttacaggctggaacatagaagggttcgatgtaccgtatattatgaatcgggtaaagatgatcctcggagagagaagcatgaaaagattttcacctattggcagagtgaaatctaagttgatacaaaacatgtatggctcaaaagagatctattcaatagatggagttagcatactcgactacctggatctgtataaaaagtttgcttttaccaacttgcctagcttctcccttgaaagtgtcgcccaacacgagaccaagaaaggtaagctgccgtacgatggcccgattaataaactgcgcgagaccaaccatcaaagatatattagctacaacataattgatgtcgaatctgtgcaagccattgataagataaggggctttatcgaccttgtcctgtcaatgtcctattacgccaaaatgccgttctcaggtgtaatgtcacccataaagacgtgggatgcgatcatcttcaattctctcaagggagagcacaaggtgatcccccaacaggggtcccacgtaaagcagtccttcccgggagcttttgtctttgagccaaagccgatcgcccgaaggtatatcatgtctttcgaccttacgtcactttacccttcaattattcgacaagtgaatatatcacccgagactatccggggccagtttaaggtacacccaatccatgagtacatagccggtacagccccaaaacccagtgacgaatactcttgcagccctaatgggtggatgtacgacaaacaccaggagggcataatcccaaaggaaatcgcgaaagtatttttccaacggaaagactggaagaaaaagatgttcgcggaggaaatgaacgccgaggctattaaaaaaatcattatgaagggagcgggtagctgttctaccaagccagaggtagagcgctacgtcaaattcagtgatgacttccttaatgagctgagtaactacacagagtctgtactgaactcactgattgaggaatgtgaaaaagccgcaacacttgctaataccaatcaactgaatcggaagatcctgattaattcactgtatggcgccttgggcaacattcatttcagatactacgacctcaggaatgccacggccattacaattttcggtcaggtcgggatccagtggatcgcccgaaaaatcaacgagtacctcaataaagtgtgtggtaccaatgacgaggattttatcgcagcaggcgataccgatagcgtgtatgtttgcgtcgacaaggtcattgaaaaggtagggctggatcggtttaaggagcagaatgatcttgtcgagtttatgaaccagtttggtaaaaaaaagatggaaccgatgatagatgtagcgtaccgagaactttgtgactacatgaataatcgcgagcacttgatgcacatggacagggaagcgatttcatgccccccactcggttcaaagggcgtagggggtttctggaaagctaagaaacggtacgccctcaacgtctatgacatggaagacaagaggttcgcggaacctcatttgaagataatggggatggagacgcaacagtcctcaactccaaaggctgtgcaagaggctctggaagaaagcatacgacgcatactccaggagggggaagagagtgttcaggagtattataaaaactttgaaaaggagtaccgccagcttgactacaaggtaatcgcggaggttaagactgcgaatgatatcgccaaatatgatgataaaggatggcccggtttcaaatgccctttccatatacgaggggtcctcacctaccgccgcgccgtgtctggtctgggggtcgcaccaattctcgacggaaataaggttatggtactcccactccgcgaggggaatccgtttggtgacaaatgcatcgcctggccgtctggtacggagctccccaaggaaatacgcagcgacgtcctcagttggatcgaccactccacactgtttcagaagtcattcgttaaacctctggccgggatgtgtgaatccgcgggtatggactacgaggagaaagcttcattggactttcttttcggtggtggttctccaaagaagaagcggaaggtctag

3xFlagtag

Cas9

40SVNLS

Nucleoplasmin NLS

P2A linker

T4DNA polymerase

SpCas9 iFE (SpCas9-GS-POLB):

atggactataaggaccacgacggagactacaaggatcatgatattgattacaaagacgatgacgataagatggccccaaagaagaagcggaaggtcggtatccacggagtcccagcagccgacaagaagtacagcatcggcctggacatcggcaccaactctgtgggctgggccgtgatcaccgacgagtacaaggtgcccagcaagaaattcaaggtgctgggcaacaccgaccggcacagcatcaagaagaacctgatcggagccctgctgttcgacagcggcgaaacagccgaggccacccggctgaagagaaccgccagaagaagatacaccagacggaagaaccggatctgctatctgcaagagatcttcagcaacgagatggccaaggtggacgacagcttcttccacagactggaagagtccttcctggtggaagaggataagaagcacgagcggcaccccatcttcggcaacatcgtggacgaggtggcctaccacgagaagtaccccaccatctaccacctgagaaagaaactggtggacagcaccgacaaggccgacctgcggctgatctatctggccctggcccacatgatcaagttccggggccacttcctgatcgagggcgacctgaaccccgacaacagcgacgttgacaagctgttcatccagctggtgcagacctacaaccagctgttcgaggaaaaccccatcaacgccagcggcgtggacgccaaggccatcctgtctgccagactgagcaagagcagacggctggaaaatctgatcgcccagctgcccggcgagaagaagaatggcctgttcggaaacctgattgccctgagcctgggcctgacccccaacttcaagagcaacttcgacctggccgaggatgccaaactgcagctgagcaaggacacctacgacgacgacctggacaacctgctggcccagatcggcgaccagtacgccgacctgtttctggccgccaagaacctgtccgacgccatcctgctgagcgacatcctgagagtgaacaccgagatcaccaaggcccccctgagcgcctctatgatcaagagatacgacgagcaccaccaggacctgaccctgctgaaagctctcgtgcggcagcagctgcctgagaagtacaaagagattttcttcgaccagagcaagaacggctacgccggctacattgacggcggagccagccaggaagagttctacaagttcatcaagcccatcctggaaaagatggacggcaccgaggaactgctcgtgaagctgaacagagaggacctgctgcggaagcagcggaccttcgacaacggcagcatcccccaccagatccacctgggagagctgcacgccattctgcggcggcaggaagatttttacccattcctgaaggacaaccgggaaaagatcgagaagatcctgaccttccgcatcccctactacgtgggccctctggccaggggaaacagcagattcgcctggatgaccagaaagagcgaggaaaccatcaccccctggaacttcgaggaagtggtggacaagggcgcttccgcccagagcttcatcgagcggatgaccaacttcgataagaacctgcccaacgagaaggtgctgcccaagcacagcctgctgtacgagtacttcaccgtgtataacgagctgaccaaagtgaaatacgtgaccgagggaatgagaaagcccgccttcctgagcggcgagcagaaaaaggccatcgtggacctgctgttcaagaccaaccggaaagtgaccgtgaagcagctgaaagaggactacttcaagaaaatcgagtgcttcgactccgtggaaatctccggcgtggaagatcggttcaacgcctccctgggcacataccacgatctgctgaaaattatcaaggacaaggacttcctggacaatgaggaaaacgaggacattctggaagatatcgtgctgaccctgacactgtttgaggacagagagatgatcgaggaacggctgaaaacctatgcccacctgttcgacgacaaagtgatgaagcagctgaagcggcggagatacaccggctggggcaggctgagccggaagctgatcaacggcatccgggacaagcagtccggcaagacaatcctggatttcctgaagtccgacggcttcgccaacagaaacttcatgcagctgatccacgacgacagcctgacctttaaagaggacatccagaaagcccaggtgtccggccagggcgatagcctgcacgagcacattgccaatctggccggcagccccgccattaagaagggcatcctgcagacagtgaaggtggtggacgagctcgtgaaagtgatgggccggcacaagcccgagaacatcgtgatcgaaatggccagagagaaccagaccacccagaagggacagaagaacagccgcgagagaatgaagcggatcgaagagggcatcaaagagctgggcagccagatcctgaaagaacaccccgtggaaaacacccagctgcagaacgagaagctgtacctgtactacctgcagaatgggcgggatatgtacgtggaccaggaactggacatcaaccggctgtccgactacgatgtggaccatatcgtgcctcagagctttctgaaggacgactccatcgacaacaaggtgctgaccagaagcgacaagaaccggggcaagagcgacaacgtgccctccgaagaggtcgtgaagaagatgaagaactactggcggcagctgctgaacgccaagctgattacccagagaaagttcgacaatctgaccaaggccgagagaggcggcctgagcgaactggataaggccggcttcatcaagagacagctggtggaaacccggcagatcacaaagcacgtggcacagatcctggactcccggatgaacactaagtacgacgagaatgacaagctgatccgggaagtgaaagtgatcaccctgaagtccaagctggtgtccgatttccggaaggatttccagttttacaaagtgcgcgagatcaacaactaccaccacgcccacgacgcctacctgaacgccgtcgtgggaaccgccctgatcaaaaagtaccctaagctggaaagcgagttcgtgtacggcgactacaaggtgtacgacgtgcggaagatgatcgccaagagcgagcaggaaatcggcaaggctaccgccaagtacttcttctacagcaacatcatgaactttttcaagaccgagattaccctggccaacggcgagatccggaagcggcctctgatcgagacaaacggcgaaaccggggagatcgtgtgggataagggccgggattttgccaccgtgcggaaagtgctgagcatgccccaagtgaatatcgtgaaaaagaccgaggtgcagacaggcggcttcagcaaagagtctatcctgcccaagaggaacagcgataagctgatcgccagaaagaaggactgggaccctaagaagtacggcggcttcgacagccccaccgtggcctattctgtgctggtggtggccaaagtggaaaagggcaagtccaagaaactgaagagtgtgaaagagctgctggggatcaccatcatggaaagaagcagcttcgagaagaatcccatcgactttctggaagccaagggctacaaagaagtgaaaaaggacctgatcatcaagctgcctaagtactccctgttcgagctggaaaacggccggaagagaatgctggcctctgccggcgaactgcagaagggaaacgaactggccctgccctccaaatatgtgaacttcctgtacctggccagccactatgagaagctgaagggctcccccgaggataatgagcagaaacagctgtttgtggaacagcacaagcactacctggacgagatcatcgagcagatcagcgagttctccaagagagtgatcctggccgacgctaatctggacaaagtgctgtccgcctacaacaagcaccgggataagcccatcagagagcaggccgagaatatcatccacctgtttaccctgaccaatctgggagcccctgccgccttcaagtactttgacaccaccatcgaccggaagaggtacaccagcaccaaagaggtgctggacgccaccctgatccaccagagcatcaccggcctgtacgagacacggatcgacctgtctcagctgggaggcgacaaaaggccggcggccacgaaaaaggccggccaggcaaaaaagaaaaagggcggcggatctggtggtgggtctggcggaggctcgtctaagcgaaaggctccgcaggagacccttaacggtggcatcacggatatgctgacggagctggctaattttgaaaaaaatgtttcacaagctatccacaagtacaacgcctacagaaaagctgcttctgtaatagccaaatatcctcataaaatcaaatcaggggccgaggccaaaaaactgccgggggtggggacgaaaatcgcagaaaaaatagatgagtttttggcgacgggtaaattgcgaaagctggaaaagatcaggcaggacgacacctcttctagtattaatttcctcacgagagtaagcggtatcggacctagtgccgccaggaagttcgtagatgaggggattaagacattggaagatctccggaagaacgaggacaagttgaatcatcaccagcgaataggacttaaatatttcggagacttcgaaaagcgcattccacgagaggagatgctgcaaatgcaagacatagtgcttaatgaagtcaaaaaagtagacagcgagtacatagctactgtctgcggatcttttaggcggggtgctgagtcttctggtgatatggatgtactgctgactcatccatcttttacaagcgagagcactaagcaaccaaaactgcttcaccaagtcgtcgagcaactgcagaaggtacactttatcactgacactctgtctaaaggtgaaacgaagttcatgggcgtctgccagttgccgagtaaaaatgatgaaaaagaatatccgcatcggagaatagacatccgactcatcccgaaggaccaatactactgtggggtcttgtacttcacgggctcagacattttcaacaagaatatgagggcacacgccctggaaaagggttttacaataaatgagtacacgatcagacctctgggagtcaccggggttgcgggagagccgcttccggttgactcagaaaaggacattttcgactatatccagtggaaatacagagaaccaaaggacagaagtgagggtggttctccaaagaagaagcggaaggtctag

3xFlagtag

Cas9

40SVNLS

Nucleoplasmin NLS

Polymerase beta

SpCas9 dFE (SpCas9-GS-4pol):

atggactataaggaccacgacggagactacaaggatcatgatattgattacaaagacgatgacgataagatggccccaaagaagaagcggaaggtcggtatccacggagtcccagcagccgacaagaagtacagcatcggcctggacatcggcaccaactctgtgggctgggccgtgatcaccgacgagtacaaggtgcccagcaagaaattcaaggtgctgggcaacaccgaccggcacagcatcaagaagaacctgatcggagccctgctgttcgacagcggcgaaacagccgaggccacccggctgaagagaaccgccagaagaagatacaccagacggaagaaccggatctgctatctgcaagagatcttcagcaacgagatggccaaggtggacgacagcttcttccacagactggaagagtccttcctggtggaagaggataagaagcacgagcggcaccccatcttcggcaacatcgtggacgaggtggcctaccacgagaagtaccccaccatctaccacctgagaaagaaactggtggacagcaccgacaaggccgacctgcggctgatctatctggccctggcccacatgatcaagttccggggccacttcctgatcgagggcgacctgaaccccgacaacagcgacgttgacaagctgttcatccagctggtgcagacctacaaccagctgttcgaggaaaaccccatcaacgccagcggcgtggacgccaaggccatcctgtctgccagactgagcaagagcagacggctggaaaatctgatcgcccagctgcccggcgagaagaagaatggcctgttcggaaacctgattgccctgagcctgggcctgacccccaacttcaagagcaacttcgacctggccgaggatgccaaactgcagctgagcaaggacacctacgacgacgacctggacaacctgctggcccagatcggcgaccagtacgccgacctgtttctggccgccaagaacctgtccgacgccatcctgctgagcgacatcctgagagtgaacaccgagatcaccaaggcccccctgagcgcctctatgatcaagagatacgacgagcaccaccaggacctgaccctgctgaaagctctcgtgcggcagcagctgcctgagaagtacaaagagattttcttcgaccagagcaagaacggctacgccggctacattgacggcggagccagccaggaagagttctacaagttcatcaagcccatcctggaaaagatggacggcaccgaggaactgctcgtgaagctgaacagagaggacctgctgcggaagcagcggaccttcgacaacggcagcatcccccaccagatccacctgggagagctgcacgccattctgcggcggcaggaagatttttacccattcctgaaggacaaccgggaaaagatcgagaagatcctgaccttccgcatcccctactacgtgggccctctggccaggggaaacagcagattcgcctggatgaccagaaagagcgaggaaaccatcaccccctggaacttcgaggaagtggtggacaagggcgcttccgcccagagcttcatcgagcggatgaccaacttcgataagaacctgcccaacgagaaggtgctgcccaagcacagcctgctgtacgagtacttcaccgtgtataacgagctgaccaaagtgaaatacgtgaccgagggaatgagaaagcccgccttcctgagcggcgagcagaaaaaggccatcgtggacctgctgttcaagaccaaccggaaagtgaccgtgaagcagctgaaagaggactacttcaagaaaatcgagtgcttcgactccgtggaaatctccggcgtggaagatcggttcaacgcctccctgggcacataccacgatctgctgaaaattatcaaggacaaggacttcctggacaatgaggaaaacgaggacattctggaagatatcgtgctgaccctgacactgtttgaggacagagagatgatcgaggaacggctgaaaacctatgcccacctgttcgacgacaaagtgatgaagcagctgaagcggcggagatacaccggctggggcaggctgagccggaagctgatcaacggcatccgggacaagcagtccggcaagacaatcctggatttcctgaagtccgacggcttcgccaacagaaacttcatgcagctgatccacgacgacagcctgacctttaaagaggacatccagaaagcccaggtgtccggccagggcgatagcctgcacgagcacattgccaatctggccggcagccccgccattaagaagggcatcctgcagacagtgaaggtggtggacgagctcgtgaaagtgatgggccggcacaagcccgagaacatcgtgatcgaaatggccagagagaaccagaccacccagaagggacagaagaacagccgcgagagaatgaagcggatcgaagagggcatcaaagagctgggcagccagatcctgaaagaacaccccgtggaaaacacccagctgcagaacgagaagctgtacctgtactacctgcagaatgggcgggatatgtacgtggaccaggaactggacatcaaccggctgtccgactacgatgtggaccatatcgtgcctcagagctttctgaaggacgactccatcgacaacaaggtgctgaccagaagcgacaagaaccggggcaagagcgacaacgtgccctccgaagaggtcgtgaagaagatgaagaactactggcggcagctgctgaacgccaagctgattacccagagaaagttcgacaatctgaccaaggccgagagaggcggcctgagcgaactggataaggccggcttcatcaagagacagctggtggaaacccggcagatcacaaagcacgtggcacagatcctggactcccggatgaacactaagtacgacgagaatgacaagctgatccgggaagtgaaagtgatcaccctgaagtccaagctggtgtccgatttccggaaggatttccagttttacaaagtgcgcgagatcaacaactaccaccacgcccacgacgcctacctgaacgccgtcgtgggaaccgccctgatcaaaaagtaccctaagctggaaagcgagttcgtgtacggcgactacaaggtgtacgacgtgcggaagatgatcgccaagagcgagcaggaaatcggcaaggctaccgccaagtacttcttctacagcaacatcatgaactttttcaagaccgagattaccctggccaacggcgagatccggaagcggcctctgatcgagacaaacggcgaaaccggggagatcgtgtgggataagggccgggattttgccaccgtgcggaaagtgctgagcatgccccaagtgaatatcgtgaaaaagaccgaggtgcagacaggcggcttcagcaaagagtctatcctgcccaagaggaacagcgataagctgatcgccagaaagaaggactgggaccctaagaagtacggcggcttcgacagccccaccgtggcctattctgtgctggtggtggccaaagtggaaaagggcaagtccaagaaactgaagagtgtgaaagagctgctggggatcaccatcatggaaagaagcagcttcgagaagaatcccatcgactttctggaagccaagggctacaaagaagtgaaaaaggacctgatcatcaagctgcctaagtactccctgttcgagctggaaaacggccggaagagaatgctggcctctgccggcgaactgcagaagggaaacgaactggccctgccctccaaatatgtgaacttcctgtacctggccagccactatgagaagctgaagggctcccccgaggataatgagcagaaacagctgtttgtggaacagcacaagcactacctggacgagatcatcgagcagatcagcgagttctccaagagagtgatcctggccgacgctaatctggacaaagtgctgtccgcctacaacaagcaccgggataagcccatcagagagcaggccgagaatatcatccacctgtttaccctgaccaatctgggagcccctgccgccttcaagtactttgacaccaccatcgaccggaagaggtacaccagcaccaaagaggtgctggacgccaccctgatccaccagagcatcaccggcctgtacgagacacggatcgacctgtctcagctgggaggcgacaaaaggccggcggccacgaaaaaggccggccaggcaaaaaagaaaaagggcggcggatctggtggtgggtctggcggaggctcgaaggaattctatattagtatcgagacagtgggaaacaacattgtagagcggtatatcgatgaaaacggcaaagaaaggactcgagaggtcgaataccttccgaccatgtttcgccactgtaaagaagaatctaagtacaaagatatttacgggaagaattgcgctccccaaaaatttccctccatgaaggatgctcgagactggatgaagcgcatggaggacataggtctcgaagcattggggatgaacgattttaagttggcttacatctccgacacttacgggtcagaaatagtgtacgataggaagttcgttcgcgtggcaaattgcgatatagaggtcactggagataagttcccggacccgatgaaggcggagtacgaaattgacgctataacacactatgactcaatcgacgaccggttctatgtatttgacctgctcaattccatgtacgggtctgtaagcaagtgggacgctaaactcgcggctaaacttgactgtgaaggaggggatgaagtacctcaggaaatcttggacagggtaatctacatgccctttgacaatgaacgagatatgcttatggagtacattaatttgtgggagcaaaagcgccccgcaatatttacaggctggaacatagaagggttcgatgtaccgtatattatgaatcgggtaaagatgatcctcggagagagaagcatgaaaagattttcacctattggcagagtgaaatctaagttgatacaaaacatgtatggctcaaaagagatctattcaatagatggagttagcatactcgactacctggatctgtataaaaagtttgcttttaccaacttgcctagcttctcccttgaaagtgtcgcccaacacgagaccaagaaaggtaagctgccgtacgatggcccgattaataaactgcgcgagaccaaccatcaaagatatattagctacaacataattgatgtcgaatctgtgcaagccattgataagataaggggctttatcgaccttgtcctgtcaatgtcctattacgccaaaatgccgttctcaggtgtaatgtcacccataaagacgtgggatgcgatcatcttcaattctctcaagggagagcacaaggtgatcccccaacaggggtcccacgtaaagcagtccttcccgggagcttttgtctttgagccaaagccgatcgcccgaaggtatatcatgtctttcgaccttacgtcactttacccttcaattattcgacaagtgaatatatcacccgagactatccggggccagtttaaggtacacccaatccatgagtacatagccggtacagccccaaaacccagtgacgaatactcttgcagccctaatgggtggatgtacgacaaacaccaggagggcataatcccaaaggaaatcgcgaaagtatttttccaacggaaagactggaagaaaaagatgttcgcggaggaaatgaacgccgaggctattaaaaaaatcattatgaagggagcgggtagctgttctaccaagccagaggtagagcgctacgtcaaattcagtgatgacttccttaatgagctgagtaactacacagagtctgtactgaactcactgattgaggaatgtgaaaaagccgcaacacttgctaataccaatcaactgaatcggaagatcctgattaattcactgtatggcgccttgggcaacattcatttcagatactacgacctcaggaatgccacggccattacaattttcggtcaggtcgggatccagtggatcgcccgaaaaatcaacgagtacctcaataaagtgtgtggtaccaatgacgaggattttatcgcagcaggcgataccgatagcgtgtatgtttgcgtcgacaaggtcattgaaaaggtagggctggatcggtttaaggagcagaatgatcttgtcgagtttatgaaccagtttggtaaaaaaaagatggaaccgatgatagatgtagcgtaccgagaactttgtgactacatgaataatcgcgagcacttgatgcacatggacagggaagcgatttcatgccccccactcggttcaaagggcgtagggggtttctggaaagctaagaaacggtacgccctcaacgtctatgacatggaagacaagaggttcgcggaacctcatttgaagataatggggatggagacgcaacagtcctcaactccaaaggctgtgcaagaggctctggaagaaagcatacgacgcatactccaggagggggaagagagtgttcaggagtattataaaaactttgaaaaggagtaccgccagcttgactacaaggtaatcgcggaggttaagactgcgaatgatatcgccaaatatgatgataaaggatggcccggtttcaaatgccctttccatatacgaggggtcctcacctaccgccgcgccgtgtctggtctgggggtcgcaccaattctcgacggaaataaggttatggtactcccactccgcgaggggaatccgtttggtgacaaatgcatcgcctggccgtctggtacggagctccccaaggaaatacgcagcgacgtcctcagttggatcgaccactccacactgtttcagaagtcattcgttaaacctctggccgggatgtgtgaatccgcgggtatggactacgaggagaaagcttcattggactttcttttcgggggtggttctccaaagaagaagcggaaggtctag

3xFlagtag

Cas9

40SVNLS

Nucleoplasmin NLS

T4DNA polymerase

HiFi iFE (HiFiCas9-GS-POLB):

atggactataaggaccacgacggagactacaaggatcatgatattgattacaaagacgatgacgataagatggccccaaagaagaagcggaaggtcggtatccacggagtcccagcagccgacaagaagtacagcatcggcctggacatcggcaccaactctgtgggctgggccgtgatcaccgacgagtacaaggtgcccagcaagaaattcaaggtgctgggcaacaccgaccggcacagcatcaagaagaacctgatcggagccctgctgttcgacagcggcgaaacagccgaggccacccggctgaagagaaccgccagaagaagatacaccagacggaagaaccggatctgctatctgcaagagatcttcagcaacgagatggccaaggtggacgacagcttcttccacagactggaagagtccttcctggtggaagaggataagaagcacgagcggcaccccatcttcggcaacatcgtggacgaggtggcctaccacgagaagtaccccaccatctaccacctgagaaagaaactggtggacagcaccgacaaggccgacctgcggctgatctatctggccctggcccacatgatcaagttccggggccacttcctgatcgagggcgacctgaaccccgacaacagcgacgttgacaagctgttcatccagctggtgcagacctacaaccagctgttcgaggaaaaccccatcaacgccagcggcgtggacgccaaggccatcctgtctgccagactgagcaagagcagacggctggaaaatctgatcgcccagctgcccggcgagaagaagaatggcctgttcggaaacctgattgccctgagcctgggcctgacccccaacttcaagagcaacttcgacctggccgaggatgccaaactgcagctgagcaaggacacctacgacgacgacctggacaacctgctggcccagatcggcgaccagtacgccgacctgtttctggccgccaagaacctgtccgacgccatcctgctgagcgacatcctgagagtgaacaccgagatcaccaaggcccccctgagcgcctctatgatcaagagatacgacgagcaccaccaggacctgaccctgctgaaagctctcgtgcggcagcagctgcctgagaagtacaaagagattttcttcgaccagagcaagaacggctacgccggctacattgacggcggagccagccaggaagagttctacaagttcatcaagcccatcctggaaaagatggacggcaccgaggaactgctcgtgaagctgaacagagaggacctgctgcggaagcagcggaccttcgacaacggcagcatcccccaccagatccacctgggagagctgcacgccattctgcggcggcaggaagatttttacccattcctgaaggacaaccgggaaaagatcgagaagatcctgaccttccgcatcccctactacgtgggccctctggccaggggaaacagcagattcgcctggatgaccagaaagagcgaggaaaccatcaccccctggaacttcgaggaagtggtggacaagggcgcttccgcccagagcttcatcgagcggatgaccaacttcgataagaacctgcccaacgagaaggtgctgcccaagcacagcctgctgtacgagtacttcaccgtgtataacgagctgaccaaagtgaaatacgtgaccgagggaatgagaaagcccgccttcctgagcggcgagcagaaaaaggccatcgtggacctgctgttcaagaccaaccggaaagtgaccgtgaagcagctgaaagaggactacttcaagaaaatcgagtgcttcgactccgtggaaatctccggcgtggaagatcggttcaacgcctccctgggcacataccacgatctgctgaaaattatcaaggacaaggacttcctggacaatgaggaaaacgaggacattctggaagatatcgtgctgaccctgacactgtttgaggacagagagatgatcgaggaacggctgaaaacctatgcccacctgttcgacgacaaagtgatgaagcagctgaagcggcggagatacaccggctggggcaggctgagccggaagctgatcaacggcatccgggacaagcagtccggcaagacaatcctggatttcctgaagtccgacggcttcgccaacgccaacttcatgcagctgatccacgacgacagcctgacctttaaagaggacatccagaaagcccaggtgtccggccagggcgatagcctgcacgagcacattgccaatctggccggcagccccgccattaagaagggcatcctgcagacagtgaaggtggtggacgagctcgtgaaagtgatgggccggcacaagcccgagaacatcgtgatcgaaatggccagagagaaccagaccacccagaagggacagaagaacagccgcgagagaatgaagcggatcgaagagggcatcaaagagctgggcagccagatcctgaaagaacaccccgtggaaaacacccagctgcagaacgagaagctgtacctgtactacctgcagaatgggcgggatatgtacgtggaccaggaactggacatcaaccggctgtccgactacgatgtggaccatatcgtgcctcagagctttctgaaggacgactccatcgacaacaaggtgctgaccagaagcgacaagaaccggggcaagagcgacaacgtgccctccgaagaggtcgtgaagaagatgaagaactactggcggcagctgctgaacgccaagctgattacccagagaaagttcgacaatctgaccaaggccgagagaggcggcctgagcgaactggataaggccggcttcatcaagagacagctggtggaaacccggcagatcacaaagcacgtggcacagatcctggactcccggatgaacactaagtacgacgagaatgacaagctgatccgggaagtgaaagtgatcaccctgaagtccaagctggtgtccgatttccggaaggatttccagttttacaaagtgcgcgagatcaacaactaccaccacgcccacgacgcctacctgaacgccgtcgtgggaaccgccctgatcaaaaagtaccctaagctggaaagcgagttcgtgtacggcgactacaaggtgtacgacgtgcggaagatgatcgccaagagcgagcaggaaatcggcaaggctaccgccaagtacttcttctacagcaacatcatgaactttttcaagaccgagattaccctggccaacggcgagatccggaagcggcctctgatcgagacaaacggcgaaaccggggagatcgtgtgggataagggccgggattttgccaccgtgcggaaagtgctgagcatgccccaagtgaatatcgtgaaaaagaccgaggtgcagacaggcggcttcagcaaagagtctatcctgcccaagaggaacagcgataagctgatcgccagaaagaaggactgggaccctaagaagtacggcggcttcgacagccccaccgtggcctattctgtgctggtggtggccaaagtggaaaagggcaagtccaagaaactgaagagtgtgaaagagctgctggggatcaccatcatggaaagaagcagcttcgagaagaatcccatcgactttctggaagccaagggctacaaagaagtgaaaaaggacctgatcatcaagctgcctaagtactccctgttcgagctggaaaacggccggaagagaatgctggcctctgccggcgaactgcagaagggaaacgaactggccctgccctccaaatatgtgaacttcctgtacctggccagccactatgagaagctgaagggctcccccgaggataatgagcagaaacagctgtttgtggaacagcacaagcactacctggacgagatcatcgagcagatcagcgagttctccaagagagtgatcctggccgacgctaatctggacaaagtgctgtccgcctacaacaagcaccgggataagcccatcagagagcaggccgagaatatcatccacctgtttaccctgaccaatctgggagcccctgccgccttcaagtactttgacaccaccatcgaccggaagaggtacaccagcaccaaagaggtgctggacgccaccctgatccaccagagcatcaccggcctgtacgagacacggatcgacctgtctcagctgggaggcgacaaaaggccggcggccacgaaaaaggccggccaggcaaaaaagaaaaagggcggcggatctggtggtgggtctggcggaggctcgtctaagcgaaaggctccgcaggagacccttaacggtggcatcacggatatgctgacggagctggctaattttgaaaaaaatgtttcacaagctatccacaagtacaacgcctacagaaaagctgcttctgtaatagccaaatatcctcataaaatcaaatcaggggccgaggccaaaaaactgccgggggtggggacgaaaatcgcagaaaaaatagatgagtttttggcgacgggtaaattgcgaaagctggaaaagatcaggcaggacgacacctcttctagtattaatttcctcacgagagtaagcggtatcggacctagtgccgccaggaagttcgtagatgaggggattaagacattggaagatctccggaagaacgaggacaagttgaatcatcaccagcgaataggacttaaatatttcggagacttcgaaaagcgcattccacgagaggagatgctgcaaatgcaagacatagtgcttaatgaagtcaaaaaagtagacagcgagtacatagctactgtctgcggatcttttaggcggggtgctgagtcttctggtgatatggatgtactgctgactcatccatcttttacaagcgagagcactaagcaaccaaaactgcttcaccaagtcgtcgagcaactgcagaaggtacactttatcactgacactctgtctaaaggtgaaacgaagttcatgggcgtctgccagttgccgagtaaaaatgatgaaaaagaatatccgcatcggagaatagacatccgactcatcccgaaggaccaatactactgtggggtcttgtacttcacgggctcagacattttcaacaagaatatgagggcacacgccctggaaaagggttttacaataaatgagtacacgatcagacctctgggagtcaccggggttgcgggagagccgcttccggttgactcagaaaaggacattttcgactatatccagtggaaatacagagaaccaaaggacagaagtgagggtggttctccaaagaagaagcggaaggtctag

3xFlagtag

Cas9 (Underbar:R691A mutation)

40SVNLS

Nucleoplasmin NLS

Polymerase beta

HiFi dFE (HiFi Cas9-GS-T4pol):

atggactataaggaccacgacggagactacaaggatcatgatattgattacaaagacgatgacgataagatggccccaaagaagaagcggaaggtcggtatccacggagtcccagcagccgacaagaagtacagcatcggcctggacatcggcaccaactctgtgggctgggccgtgatcaccgacgagtacaaggtgcccagcaagaaattcaaggtgctgggcaacaccgaccggcacagcatcaagaagaacctgatcggagccctgctgttcgacagcggcgaaacagccgaggccacccggctgaagagaaccgccagaagaagatacaccagacggaagaaccggatctgctatctgcaagagatcttcagcaacgagatggccaaggtggacgacagcttcttccacagactggaagagtccttcctggtggaagaggataagaagcacgagcggcaccccatcttcggcaacatcgtggacgaggtggcctaccacgagaagtaccccaccatctaccacctgagaaagaaactggtggacagcaccgacaaggccgacctgcggctgatctatctggccctggcccacatgatcaagttccggggccacttcctgatcgagggcgacctgaaccccgacaacagcgacgttgacaagctgttcatccagctggtgcagacctacaaccagctgttcgaggaaaaccccatcaacgccagcggcgtggacgccaaggccatcctgtctgccagactgagcaagagcagacggctggaaaatctgatcgcccagctgcccggcgagaagaagaatggcctgttcggaaacctgattgccctgagcctgggcctgacccccaacttcaagagcaacttcgacctggccgaggatgccaaactgcagctgagcaaggacacctacgacgacgacctggacaacctgctggcccagatcggcgaccagtacgccgacctgtttctggccgccaagaacctgtccgacgccatcctgctgagcgacatcctgagagtgaacaccgagatcaccaaggcccccctgagcgcctctatgatcaagagatacgacgagcaccaccaggacctgaccctgctgaaagctctcgtgcggcagcagctgcctgagaagtacaaagagattttcttcgaccagagcaagaacggctacgccggctacattgacggcggagccagccaggaagagttctacaagttcatcaagcccatcctggaaaagatggacggcaccgaggaactgctcgtgaagctgaacagagaggacctgctgcggaagcagcggaccttcgacaacggcagcatcccccaccagatccacctgggagagctgcacgccattctgcggcggcaggaagatttttacccattcctgaaggacaaccgggaaaagatcgagaagatcctgaccttccgcatcccctactacgtgggccctctggccaggggaaacagcagattcgcctggatgaccagaaagagcgaggaaaccatcaccccctggaacttcgaggaagtggtggacaagggcgcttccgcccagagcttcatcgagcggatgaccaacttcgataagaacctgcccaacgagaaggtgctgcccaagcacagcctgctgtacgagtacttcaccgtgtataacgagctgaccaaagtgaaatacgtgaccgagggaatgagaaagcccgccttcctgagcggcgagcagaaaaaggccatcgtggacctgctgttcaagaccaaccggaaagtgaccgtgaagcagctgaaagaggactacttcaagaaaatcgagtgcttcgactccgtggaaatctccggcgtggaagatcggttcaacgcctccctgggcacataccacgatctgctgaaaattatcaaggacaaggacttcctggacaatgaggaaaacgaggacattctggaagatatcgtgctgaccctgacactgtttgaggacagagagatgatcgaggaacggctgaaaacctatgcccacctgttcgacgacaaagtgatgaagcagctgaagcggcggagatacaccggctggggcaggctgagccggaagctgatcaacggcatccgggacaagcagtccggcaagacaatcctggatttcctgaagtccgacggcttcgccaacgccaacttcatgcagctgatccacgacgacagcctgacctttaaagaggacatccagaaagcccaggtgtccggccagggcgatagcctgcacgagcacattgccaatctggccggcagccccgccattaagaagggcatcctgcagacagtgaaggtggtggacgagctcgtgaaagtgatgggccggcacaagcccgagaacatcgtgatcgaaatggccagagagaaccagaccacccagaagggacagaagaacagccgcgagagaatgaagcggatcgaagagggcatcaaagagctgggcagccagatcctgaaagaacaccccgtggaaaacacccagctgcagaacgagaagctgtacctgtactacctgcagaatgggcgggatatgtacgtggaccaggaactggacatcaaccggctgtccgactacgatgtggaccatatcgtgcctcagagctttctgaaggacgactccatcgacaacaaggtgctgaccagaagcgacaagaaccggggcaagagcgacaacgtgccctccgaagaggtcgtgaagaagatgaagaactactggcggcagctgctgaacgccaagctgattacccagagaaagttcgacaatctgaccaaggccgagagaggcggcctgagcgaactggataaggccggcttcatcaagagacagctggtggaaacccggcagatcacaaagcacgtggcacagatcctggactcccggatgaacactaagtacgacgagaatgacaagctgatccgggaagtgaaagtgatcaccctgaagtccaagctggtgtccgatttccggaaggatttccagttttacaaagtgcgcgagatcaacaactaccaccacgcccacgacgcctacctgaacgccgtcgtgggaaccgccctgatcaaaaagtaccctaagctggaaagcgagttcgtgtacggcgactacaaggtgtacgacgtgcggaagatgatcgccaagagcgagcaggaaatcggcaaggctaccgccaagtacttcttctacagcaacatcatgaactttttcaagaccgagattaccctggccaacggcgagatccggaagcggcctctgatcgagacaaacggcgaaaccggggagatcgtgtgggataagggccgggattttgccaccgtgcggaaagtgctgagcatgccccaagtgaatatcgtgaaaaagaccgaggtgcagacaggcggcttcagcaaagagtctatcctgcccaagaggaacagcgataagctgatcgccagaaagaaggactgggaccctaagaagtacggcggcttcgacagccccaccgtggcctattctgtgctggtggtggccaaagtggaaaagggcaagtccaagaaactgaagagtgtgaaagagctgctggggatcaccatcatggaaagaagcagcttcgagaagaatcccatcgactttctggaagccaagggctacaaagaagtgaaaaaggacctgatcatcaagctgcctaagtactccctgttcgagctggaaaacggccggaagagaatgctggcctctgccggcgaactgcagaagggaaacgaactggccctgccctccaaatatgtgaacttcctgtacctggccagccactatgagaagctgaagggctcccccgaggataatgagcagaaacagctgtttgtggaacagcacaagcactacctggacgagatcatcgagcagatcagcgagttctccaagagagtgatcctggccgacgctaatctggacaaagtgctgtccgcctacaacaagcaccgggataagcccatcagagagcaggccgagaatatcatccacctgtttaccctgaccaatctgggagcccctgccgccttcaagtactttgacaccaccatcgaccggaagaggtacaccagcaccaaagaggtgctggacgccaccctgatccaccagagcatcaccggcctgtacgagacacggatcgacctgtctcagctgggaggcgacaaaaggccggcggccacgaaaaaggccggccaggcaaaaaagaaaaagggcggcggatctggtggtgggtctggcggaggctcgaaggaattctatattagtatcgagacagtgggaaacaacattgtagagcggtatatcgatgaaaacggcaaagaaaggactcgagaggtcgaataccttccgaccatgtttcgccactgtaaagaagaatctaagtacaaagatatttacgggaagaattgcgctccccaaaaatttccctccatgaaggatgctcgagactggatgaagcgcatggaggacataggtctcgaagcattggggatgaacgattttaagttggcttacatctccgacacttacgggtcagaaatagtgtacgataggaagttcgttcgcgtggcaaattgcgatatagaggtcactggagataagttcccggacccgatgaaggcggagtacgaaattgacgctataacacactatgactcaatcgacgaccggttctatgtatttgacctgctcaattccatgtacgggtctgtaagcaagtgggacgctaaactcgcggctaaacttgactgtgaaggaggggatgaagtacctcaggaaatcttggacagggtaatctacatgccctttgacaatgaacgagatatgcttatggagtacattaatttgtgggagcaaaagcgccccgcaatatttacaggctggaacatagaagggttcgatgtaccgtatattatgaatcgggtaaagatgatcctcggagagagaagcatgaaaagattttcacctattggcagagtgaaatctaagttgatacaaaacatgtatggctcaaaagagatctattcaatagatggagttagcatactcgactacctggatctgtataaaaagtttgcttttaccaacttgcctagcttctcccttgaaagtgtcgcccaacacgagaccaagaaaggtaagctgccgtacgatggcccgattaataaactgcgcgagaccaaccatcaaagatatattagctacaacataattgatgtcgaatctgtgcaagccattgataagataaggggctttatcgaccttgtcctgtcaatgtcctattacgccaaaatgccgttctcaggtgtaatgtcacccataaagacgtgggatgcgatcatcttcaattctctcaagggagagcacaaggtgatcccccaacaggggtcccacgtaaagcagtccttcccgggagcttttgtctttgagccaaagccgatcgcccgaaggtatatcatgtctttcgaccttacgtcactttacccttcaattattcgacaagtgaatatatcacccgagactatccggggccagtttaaggtacacccaatccatgagtacatagccggtacagccccaaaacccagtgacgaatactcttgcagccctaatgggtggatgtacgacaaacaccaggagggcataatcccaaaggaaatcgcgaaagtatttttccaacggaaagactggaagaaaaagatgttcgcggaggaaatgaacgccgaggctattaaaaaaatcattatgaagggagcgggtagctgttctaccaagccagaggtagagcgctacgtcaaattcagtgatgacttccttaatgagctgagtaactacacagagtctgtactgaactcactgattgaggaatgtgaaaaagccgcaacacttgctaataccaatcaactgaatcggaagatcctgattaattcactgtatggcgccttgggcaacattcatttcagatactacgacctcaggaatgccacggccattacaattttcggtcaggtcgggatccagtggatcgcccgaaaaatcaacgagtacctcaataaagtgtgtggtaccaatgacgaggattttatcgcagcaggcgataccgatagcgtgtatgtttgcgtcgacaaggtcattgaaaaggtagggctggatcggtttaaggagcagaatgatcttgtcgagtttatgaaccagtttggtaaaaaaaagatggaaccgatgatagatgtagcgtaccgagaactttgtgactacatgaataatcgcgagcacttgatgcacatggacagggaagcgatttcatgccccccactcggttcaaagggcgtagggggtttctggaaagctaagaaacggtacgccctcaacgtctatgacatggaagacaagaggttcgcggaacctcatttgaagataatggggatggagacgcaacagtcctcaactccaaaggctgtgcaagaggctctggaagaaagcatacgacgcatactccaggagggggaagagagtgttcaggagtattataaaaactttgaaaaggagtaccgccagcttgactacaaggtaatcgcggaggttaagactgcgaatgatatcgccaaatatgatgataaaggatggcccggtttcaaatgccctttccatatacgaggggtcctcacctaccgccgcgccgtgtctggtctgggggtcgcaccaattctcgacggaaataaggttatggtactcccactccgcgaggggaatccgtttggtgacaaatgcatcgcctggccgtctggtacggagctccccaaggaaatacgcagcgacgtcctcagttggatcgaccactccacactgtttcagaagtcattcgttaaacctctggccgggatgtgtgaatccgcgggtatggactacgaggagaaagcttcattggactttcttttcgggggtggttctccaaagaagaagcggaaggtctag

R691A mutation

3xFlagtag

Cas9 (Underbar:R691A mutation)

40SVNLS

Nucleoplasmin NLS

T4DNA polymerase

LZ3 iFE (LZ3 Cas9-GS-POLB):

atggactataaggaccacgacggagactacaaggatcatgatattgattacaaagacgatgacgataagatggccccaaagaagaagcggaaggtcggtatccacggagtcccagcagccgacaagaagtacagcatcggcctggacatcggcaccaactctgtgggctgggccgtgatcaccgacgagtacaaggtgcccagcaagaaattcaaggtgctgggcaacaccgaccggcacagcatcaagaagaacctgatcggagccctgctgttcgacagcggcgaaacagccgaggccacccggctgaagagaaccgccagaagaagatacaccagacggaagaaccggatctgctatctgcaagagatcttcagcaacgagatggccaaggtggacgacagcttcttccacagactggaagagtccttcctggtggaagaggataagaagcacgagcggcaccccatcttcggcaacatcgtggacgaggtggcctaccacgagaagtaccccaccatctaccacctgagaaagaaactggtggacagcaccgacaaggccgacctgcggctgatctatctggccctggcccacatgatcaagttccggggccacttcctgatcgagggcgacctgaaccccgacaacagcgacgtggacaagctgttcatccagctggtgcagacctacaaccagctgttcgaggaaaaccccatcaacgccagcggcgtggacgccaaggccatcctgtctgccagactgagcaagagcagacggctggaaaatctgatcgcccagctgcccggcgagaagaagaatggcctgttcggaaacctgattgccctgagcctgggcctgacccccaacttcaagagcaacttcgacctggccgaggatgccaaactgcagctgagcaaggacacctacgacgacgacctggacaacctgctggcccagatcggcgaccagtacgccgacctgtttctggccgccaagaacctgtccgacgccatcctgctgagcgacatcctgagagtgaacaccgagatcaccaaggcccccctgagcgcctctatgatcaagagatacgacgagcaccaccaggacctgaccctgctgaaagctctcgtgcggcagcagctgcctgagaagtacaaagagattttcttcgaccagagcaagaacggctacgccggctacattgacggcggagccagccaggaagagttctacaagttcatcaagcccatcctggaaaagatggacggcaccgaggaactgctcgtgaagctgaacagagaggacctgctgcggaagcagcggaccttcgacaacggcagcatcccccaccagatccacctgggagagctgcacgccattctgcggcggcaggaagatttttacccattcctgaaggacaaccgggaaaagatcgagaagatcctgaccttccgcatcccctactacgtgggccctctggccaggggaaacagcagattcgcctggatgaccagaaagagcgaggaaaccatcaccccctggaacttcgaggaagtggtggacaagggcgcttccgcccagagcttcatcgagcggatgaccaacttcgataagaacctgcccaacgagaaggtgctgcccaagcacagcctgctgtacgagtacttcaccgtgtataacgagctgaccaaagtgaaatacgtgaccgagggaatgagaaagcccgccttcctgagcggcgagcagaaaaaggccatcgtggacctgctgttcaagaccaaccggaaagtgaccgtgaagcagctgaaagaggactacttcaagaaaatcgagtgcttcgactccgtggaaatctccggcgtggaagatcggttcaacgcctccctgggcacataccacgatctgctgaaaattatcaaggacaaggacttcctggacaatgaggaaaacgaggacattctggaagatatcgtgctgaccctgacactgtttgaggacagagagatgatcgaggaacggctgaaaacctatgcccacctgttcgacgacaaagtgatgaagcagctgaagcggcggagatacaccggctggggcaggctgagccggaagctgatcaacggcatccgggacaagcagtccggcaagacaatcctggatttcctgaagtccgacggcttcgcctgcagaaacttcatgcagctgatccacgacgacagcctgacctttaaagaggacatccagaaagcccaggtgtccggccagggcgatagcctgcacgagcacattgccaatctggccggcagccccgccattaagaagggcatcctgcagacagtgaaggtggtggacgagctcgtgaaagtgatgggccggcacaagcccgagaacatcgtgatcgaaatggccagagagaaccagatcacccagaagggacagaagaacagccgcgagagaatgaagcggatcgaagagggcatcaaagagctgggcagccagatcctgaaagaacaccccgtggaaaacacccagctgcagaacgagaagctgtacctgtactacctgcagaatgggcgggatatgtacgtggaccaggaactggacatcaaccggctgtccgactacgatgtggaccatatcgtgcctcagagctttctgaaggacgactccatcgacaacaaggtgctgaccagaagcgacaagaaccggggcaagagcgacaacgtgccctccgaagaggtcgtgaagaagatgaagaactactggcggcagctgctgaacgccaagctgattacccagagaaagttcgacaatctgaccaaggccgagagaggcggcctgagcgaactggataaggccatgttcatcaagagacagctggtggaaacccggcagatcacaaagcacgtggcacagatcctggactcccggatgaacactaagtacgacgagaatgacaagctgatccgggaagtgaaagtgatcaccctgaagtccaagctggtgtccgatttccggaaggatttccagttttacaaagtgcgcgagatcaacaaataccaccacgcccacgacgcctacctgaacgccgtcgtgggaaccgccctgatcaaaaagtaccctaagctggaaagcgagttcgtgtacggcgactacaaggtgtacgacgtgcggaagatgatcgccaagagcgagcaggaaatcggcaaggctaccgccaagtacttcttctacagcaacatcatgaactttttcaagaccgagattaccctggccaacggcgagatccggaagcggcctctgatcgagacaaacggcgaaaccggggagatcgtgtgggataagggccgggattttgccaccgtgcggaaagtgctgagcatgccccaagtgaatatcgtgaaaaagaccgaggtgcagacaggcggcttcagcaaagagtctatcctgcccaagaggaacagcgataagctgatcgccagaaagaaggactgggaccctaagaagtacggcggcttcgacagccccaccgtggcctattctgtgctggtggtggccaaagtggaaaagggcaagtccaagaaactgaagagtgtgaaagagctgctggggatcaccatcatggaaagaagcagcttcgagaagaatcccatcgactttctggaagccaagggctacaaagaagtgaaaaaggacctgatcatcaagctgcctaagtactccctgttcgagctggaaaacggccggaagagaatgctggcctctgccggcgaactgcagaagggaaacgaactggccctgccctccaaatatgtgaacttcctgtacctggccagccactatgagaagctgaagggctcccccgaggataatgagcagaaacagctgtttgtggaacagcacaagcactacctggacgagatcatcgagcagatcagcgagttctccaagagagtgatcctggccgacgctaatctggacaaagtgctgtccgcctacaacaagcaccgggataagcccatcagagagcaggccgagaatatcatccacctgtttaccctgaccaatctgggagcccctgccgccttcaagtactttgacaccaccatcgaccggaagaggtacaccagcaccaaagaggtgctggacgccaccctgatccaccagagcatcaccggcctgtacgagacacggatcgacctgtctcagctgggaggcgacaaaaggccggcggccacgaaaaaggccggccaggcaaaaaagaaaaagggcggcggatctggtggtgggtctggcggaggctcgtctaagcgaaaggctccgcaggagacccttaacggtggcatcacggatatgctgacggagctggctaattttgaaaaaaatgtttcacaagctatccacaagtacaacgcctacagaaaagctgcttctgtaatagccaaatatcctcataaaatcaaatcaggggccgaggccaaaaaactgccgggggtggggacgaaaatcgcagaaaaaatagatgagtttttggcgacgggtaaattgcgaaagctggaaaagatcaggcaggacgacacctcttctagtattaatttcctcacgagagtaagcggtatcggacctagtgccgccaggaagttcgtagatgaggggattaagacattggaagatctccggaagaacgaggacaagttgaatcatcaccagcgaataggacttaaatatttcggagacttcgaaaagcgcattccacgagaggagatgctgcaaatgcaagacatagtgcttaatgaagtcaaaaaagtagacagcgagtacatagctactgtctgcggatcttttaggcggggtgctgagtcttctggtgatatggatgtactgctgactcatccatcttttacaagcgagagcactaagcaaccaaaactgcttcaccaagtcgtcgagcaactgcagaaggtacactttatcactgacactctgtctaaaggtgaaacgaagttcatgggcgtctgccagttgccgagtaaaaatgatgaaaaagaatatccgcatcggagaatagacatccgactcatcccgaaggaccaatactactgtggggtcttgtacttcacgggctcagacattttcaacaagaatatgagggcacacgccctggaaaagggttttacaataaatgagtacacgatcagacctctgggagtcaccggggttgcgggagagccgcttccggttgactcagaaaaggacattttcgactatatccagtggaaatacagagaaccaaaggacagaagtgagggtggttctccaaagaagaagcggaaggtctag

3xFlagtag

LZ3Cas9

40SVNLS

Nucleoplasmin NLS

Polymerase beta

LZ3 dFE (LZ3 Cas9-GS-T4pol):

atggactataaggaccacgacggagactacaaggatcatgatattgattacaaagacgatgacgataagatggccccaaagaagaagcggaaggtcggtatccacggagtcccagcagccgacaagaagtacagcatcggcctggacatcggcaccaactctgtgggctgggccgtgatcaccgacgagtacaaggtgcccagcaagaaattcaaggtgctgggcaacaccgaccggcacagcatcaagaagaacctgatcggagccctgctgttcgacagcggcgaaacagccgaggccacccggctgaagagaaccgccagaagaagatacaccagacggaagaaccggatctgctatctgcaagagatcttcagcaacgagatggccaaggtggacgacagcttcttccacagactggaagagtccttcctggtggaagaggataagaagcacgagcggcaccccatcttcggcaacatcgtggacgaggtggcctaccacgagaagtaccccaccatctaccacctgagaaagaaactggtggacagcaccgacaaggccgacctgcggctgatctatctggccctggcccacatgatcaagttccggggccacttcctgatcgagggcgacctgaaccccgacaacagcgacgtggacaagctgttcatccagctggtgcagacctacaaccagctgttcgaggaaaaccccatcaacgccagcggcgtggacgccaaggccatcctgtctgccagactgagcaagagcagacggctggaaaatctgatcgcccagctgcccggcgagaagaagaatggcctgttcggaaacctgattgccctgagcctgggcctgacccccaacttcaagagcaacttcgacctggccgaggatgccaaactgcagctgagcaaggacacctacgacgacgacctggacaacctgctggcccagatcggcgaccagtacgccgacctgtttctggccgccaagaacctgtccgacgccatcctgctgagcgacatcctgagagtgaacaccgagatcaccaaggcccccctgagcgcctctatgatcaagagatacgacgagcaccaccaggacctgaccctgctgaaagctctcgtgcggcagcagctgcctgagaagtacaaagagattttcttcgaccagagcaagaacggctacgccggctacattgacggcggagccagccaggaagagttctacaagttcatcaagcccatcctggaaaagatggacggcaccgaggaactgctcgtgaagctgaacagagaggacctgctgcggaagcagcggaccttcgacaacggcagcatcccccaccagatccacctgggagagctgcacgccattctgcggcggcaggaagatttttacccattcctgaaggacaaccgggaaaagatcgagaagatcctgaccttccgcatcccctactacgtgggccctctggccaggggaaacagcagattcgcctggatgaccagaaagagcgaggaaaccatcaccccctggaacttcgaggaagtggtggacaagggcgcttccgcccagagcttcatcgagcggatgaccaacttcgataagaacctgcccaacgagaaggtgctgcccaagcacagcctgctgtacgagtacttcaccgtgtataacgagctgaccaaagtgaaatacgtgaccgagggaatgagaaagcccgccttcctgagcggcgagcagaaaaaggccatcgtggacctgctgttcaagaccaaccggaaagtgaccgtgaagcagctgaaagaggactacttcaagaaaatcgagtgcttcgactccgtggaaatctccggcgtggaagatcggttcaacgcctccctgggcacataccacgatctgctgaaaattatcaaggacaaggacttcctggacaatgaggaaaacgaggacattctggaagatatcgtgctgaccctgacactgtttgaggacagagagatgatcgaggaacggctgaaaacctatgcccacctgttcgacgacaaagtgatgaagcagctgaagcggcggagatacaccggctggggcaggctgagccggaagctgatcaacggcatccgggacaagcagtccggcaagacaatcctggatttcctgaagtccgacggcttcgcctgcagaaacttcatgcagctgatccacgacgacagcctgacctttaaagaggacatccagaaagcccaggtgtccggccagggcgatagcctgcacgagcacattgccaatctggccggcagccccgccattaagaagggcatcctgcagacagtgaaggtggtggacgagctcgtgaaagtgatgggccggcacaagcccgagaacatcgtgatcgaaatggccagagagaaccagatcacccagaagggacagaagaacagccgcgagagaatgaagcggatcgaagagggcatcaaagagctgggcagccagatcctgaaagaacaccccgtggaaaacacccagctgcagaacgagaagctgtacctgtactacctgcagaatgggcgggatatgtacgtggaccaggaactggacatcaaccggctgtccgactacgatgtggaccatatcgtgcctcagagctttctgaaggacgactccatcgacaacaaggtgctgaccagaagcgacaagaaccggggcaagagcgacaacgtgccctccgaagaggtcgtgaagaagatgaagaactactggcggcagctgctgaacgccaagctgattacccagagaaagttcgacaatctgaccaaggccgagagaggcggcctgagcgaactggataaggccatgttcatcaagagacagctggtggaaacccggcagatcacaaagcacgtggcacagatcctggactcccggatgaacactaagtacgacgagaatgacaagctgatccgggaagtgaaagtgatcaccctgaagtccaagctggtgtccgatttccggaaggatttccagttttacaaagtgcgcgagatcaacaaataccaccacgcccacgacgcctacctgaacgccgtcgtgggaaccgccctgatcaaaaagtaccctaagctggaaagcgagttcgtgtacggcgactacaaggtgtacgacgtgcggaagatgatcgccaagagcgagcaggaaatcggcaaggctaccgccaagtacttcttctacagcaacatcatgaactttttcaagaccgagattaccctggccaacggcgagatccggaagcggcctctgatcgagacaaacggcgaaaccggggagatcgtgtgggataagggccgggattttgccaccgtgcggaaagtgctgagcatgccccaagtgaatatcgtgaaaaagaccgaggtgcagacaggcggcttcagcaaagagtctatcctgcccaagaggaacagcgataagctgatcgccagaaagaaggactgggaccctaagaagtacggcggcttcgacagccccaccgtggcctattctgtgctggtggtggccaaagtggaaaagggcaagtccaagaaactgaagagtgtgaaagagctgctggggatcaccatcatggaaagaagcagcttcgagaagaatcccatcgactttctggaagccaagggctacaaagaagtgaaaaaggacctgatcatcaagctgcctaagtactccctgttcgagctggaaaacggccggaagagaatgctggcctctgccggcgaactgcagaagggaaacgaactggccctgccctccaaatatgtgaacttcctgtacctggccagccactatgagaagctgaagggctcccccgaggataatgagcagaaacagctgtttgtggaacagcacaagcactacctggacgagatcatcgagcagatcagcgagttctccaagagagtgatcctggccgacgctaatctggacaaagtgctgtccgcctacaacaagcaccgggataagcccatcagagagcaggccgagaatatcatccacctgtttaccctgaccaatctgggagcccctgccgccttcaagtactttgacaccaccatcgaccggaagaggtacaccagcaccaaagaggtgctggacgccaccctgatccaccagagcatcaccggcctgtacgagacacggatcgacctgtctcagctgggaggcgacaaaaggccggcggccacgaaaaaggccggccaggcaaaaaagaaaaagggcggcggatctggtggtgggtctggcggaggctcgaaggaattctatattagtatcgagacagtgggaaacaacattgtagagcggtatatcgatgaaaacggcaaagaaaggactcgagaggtcgaataccttccgaccatgtttcgccactgtaaagaagaatctaagtacaaagatatttacgggaagaattgcgctccccaaaaatttccctccatgaaggatgctcgagactggatgaagcgcatggaggacataggtctcgaagcattggggatgaacgattttaagttggcttacatctccgacacttacgggtcagaaatagtgtacgataggaagttcgttcgcgtggcaaattgcgatatagaggtcactggagataagttcccggacccgatgaaggcggagtacgaaattgacgctataacacactatgactcaatcgacgaccggttctatgtatttgacctgctcaattccatgtacgggtctgtaagcaagtgggacgctaaactcgcggctaaacttgactgtgaaggaggggatgaagtacctcaggaaatcttggacagggtaatctacatgccctttgacaatgaacgagatatgcttatggagtacattaatttgtgggagcaaaagcgccccgcaatatttacaggctggaacatagaagggttcgatgtaccgtatattatgaatcgggtaaagatgatcctcggagagagaagcatgaaaagattttcacctattggcagagtgaaatctaagttgatacaaaacatgtatggctcaaaagagatctattcaatagatggagttagcatactcgactacctggatctgtataaaaagtttgcttttaccaacttgcctagcttctcccttgaaagtgtcgcccaacacgagaccaagaaaggtaagctgccgtacgatggcccgattaataaactgcgcgagaccaaccatcaaagatatattagctacaacataattgatgtcgaatctgtgcaagccattgataagataaggggctttatcgaccttgtcctgtcaatgtcctattacgccaaaatgccgttctcaggtgtaatgtcacccataaagacgtgggatgcgatcatcttcaattctctcaagggagagcacaaggtgatcccccaacaggggtcccacgtaaagcagtccttcccgggagcttttgtctttgagccaaagccgatcgcccgaaggtatatcatgtctttcgaccttacgtcactttacccttcaattattcgacaagtgaatatatcacccgagactatccggggccagtttaaggtacacccaatccatgagtacatagccggtacagccccaaaacccagtgacgaatactcttgcagccctaatgggtggatgtacgacaaacaccaggagggcataatcccaaaggaaatcgcgaaagtatttttccaacggaaagactggaagaaaaagatgttcgcggaggaaatgaacgccgaggctattaaaaaaatcattatgaagggagcgggtagctgttctaccaagccagaggtagagcgctacgtcaaattcagtgatgacttccttaatgagctgagtaactacacagagtctgtactgaactcactgattgaggaatgtgaaaaagccgcaacacttgctaataccaatcaactgaatcggaagatcctgattaattcactgtatggcgccttgggcaacattcatttcagatactacgacctcaggaatgccacggccattacaattttcggtcaggtcgggatccagtggatcgcccgaaaaatcaacgagtacctcaataaagtgtgtggtaccaatgacgaggattttatcgcagcaggcgataccgatagcgtgtatgtttgcgtcgacaaggtcattgaaaaggtagggctggatcggtttaaggagcagaatgatcttgtcgagtttatgaaccagtttggtaaaaaaaagatggaaccgatgatagatgtagcgtaccgagaactttgtgactacatgaataatcgcgagcacttgatgcacatggacagggaagcgatttcatgccccccactcggttcaaagggcgtagggggtttctggaaagctaagaaacggtacgccctcaacgtctatgacatggaagacaagaggttcgcggaacctcatttgaagataatggggatggagacgcaacagtcctcaactccaaaggctgtgcaagaggctctggaagaaagcatacgacgcatactccaggagggggaagagagtgttcaggagtattataaaaactttgaaaaggagtaccgccagcttgactacaaggtaatcgcggaggttaagactgcgaatgatatcgccaaatatgatgataaaggatggcccggtttcaaatgccctttccatatacgaggggtcctcacctaccgccgcgccgtgtctggtctgggggtcgcaccaattctcgacggaaataaggttatggtactcccactccgcgaggggaatccgtttggtgacaaatgcatcgcctggccgtctggtacggagctccccaaggaaatacgcagcgacgtcctcagttggatcgaccactccacactgtttcagaagtcattcgttaaacctctggccgggatgtgtgaatccgcgggtatggactacgaggagaaagcttcattggactttcttttcgggggtggttctccaaagaagaagcggaaggtctag

3xFlagtag

LZ3Cas9

40SVNLS

Nucleoplasmin NLS

T4DNA polymerase

**Supplementary Sequences 2.**

ssODN for generating HEK293T cells containing RAI1^3103insC^ mutation

ATGCAGGGCACCAGTGCTGCCCAAAGACCTCTTGCTCCCTGAATCCTGCACAGGGCCCCCCCCAGGGACAGATGGAAGGGGCTGGAGCCCCAGGCCGGGGGGCCTCGGAAGGGCTCCCCA
